# Discovering the drivers of clonal hematopoiesis

**DOI:** 10.1101/2020.10.22.350140

**Authors:** Oriol Pich, Iker Reyes-Salazar, Abel Gonzalez-Perez, Nuria Lopez-Bigas

**Author notes:** Corresponding authors Abel Gonzalez-Perez, Nuria Lopez-Bigas.

## Abstract

Mutations in genes that confer a selective advantage to hematopoietic stem cells (HSCs) in certain conditions drive clonal hematopoiesis (CH). While some CH drivers have been identified experimentally or through epidemiological studies, the compendium of all genes able to drive CH upon mutations in HSCs is far from complete. We propose that identifying signals of positive selection in blood somatic mutations may be an effective way to identify CH driver genes, similarly as done to identify cancer genes. Using a reverse somatic variant calling approach, we repurposed whole-genome and whole-exome blood/tumor paired samples of more than 12,000 donors from two large cancer genomics cohorts to identify blood somatic mutations. The application of IntOGen, a robust driver discovery pipeline, to blood somatic mutations across both cohorts, and more than 24,000 targeted sequenced samples yielded a list of close to 70 genes with signals of positive selection in CH, available at http://www.intogen.org/ch. This approach recovers all known CH genes, and discovers novel candidates. Generating this compendium is an essential step to understand the molecular mechanisms of CH and to accurately detect individuals with CH to ascertain their risk to develop related diseases.

## Introduction

Clonal hematopoiesis (CH) is a condition related to aging across the human population^1–9^. It is driven by somatic alterations that appear in hematopoietic stem cells (HSCs) along their life and confer them a selective advantage. It was first recognized through cytogenetic studies in the 1960s and its genetic bases and prevalence with aging were first discovered through studies of non-random X chromosome inactivation in women^10,11^. In the past decade, genomic studies of thousands of donors without any hematologic phenotype identified causal somatic variants in genes known to be associated to hematopoietic malignancies, such as DNMT3A, TET2, ASXL1, TP53, JAK2 and SF3B1^1,2,9,10,12–15^. The progeny of the HSC bearing mutations of one of these genes enlarges over time with respect to that of other HSCs, thus occupying an increasing fraction of the hematopoietic compartment along the life of an individual. It thus develops in a process of clonal expansion. CH is known to be associated with other health risks, such as hematopoietic malignancies or increased incidence of cardiovascular disease^2,4,5,8,16^.

The aforementioned human genomic studies, and more recent analyses^16–22^ have identified a list of CH-causing somatic variants. Nevertheless, their identification is hampered by the fact that the clonal expansion related to CH is rather modest, and therefore, it appears within a range of low variant allele frequency (VAF). This peculiarity has determined the development of two main strategies of detection. On the one hand, some projects with access to very deep sequencing data of particular sites of the human genome (e.g., tumor panel sequencing) are able to identify CH with exquisite sensitivity, but only if the causing variant overlaps with the sites sequenced^14,16–18,20^. On the other, whole-genome or whole-exome sequencing data has been exploited to identify blood somatic variants at VAF above the limit of detection of bulk sequencing, but below that corresponding to germline variants^13,21–23^. This approach is thus only able to detect relatively large CH clones. One important caveat of these two types of approaches is that not all genes affected by mutations across blood samples (even known cancer driver genes) are drivers of CH. Whereas sequencing more blood samples will conceivably lead to the identification of more recurrently mutated suspicious genes, many of them are prone to be passengers of this clonal expansion process.

As a result, an accurate and complete list of CH-related genes remains elusive to date. Completing this catalog is essential to comprehensively identifying CH in individuals to accurately ascertain their risk to develop related diseases and to complete our knowledge of the molecular mechanisms underlying CH.

In recent years efforts to identify genes with mutations under positive selection in tumorigenesis have begun to uncover the compendium of mutational cancer driver genes^24–27^. We reasoned that the clonal expansion that drives clonal hematopoiesis is reminiscent of that observed in tumors, and therefore methods to detect positive selection in the mutations of genes across tumors could be applied to identify the complete list of CH-related genes. The identification of signals of positive selection across genes is expected to produce a list of CH genes with higher specificity and sensitivity than a list of recurrently mutated suspicious genes. To detect these signals of positive selection, we first need to accurately identify whole-genome blood somatic mutations. Here, we repurposed blood and tumor samples of donors with no known hematopoietic malignancy in the context of primary^28^ (N∼8,000) and metastatic^29^ (N∼4,000) cancer genomics initiatives to detect somatic mutations in blood. To this end, we used the paired tumor sample as the reference germline genome of the donors in these two cohorts. On the set of blood somatic mutations identified, we then ran the Integrative OncoGenomics (IntOGen^25^) pipeline that implements seven state-of-the-art driver discovery methods to identify signals of positive selection in the observed pattern of mutations in genes across samples. As a result, known CH-related genes and interesting novel candidates were identified. Our results open up the opportunity to repurpose cancer genomics data in the public domain to identify the compendium of CH-related genes and variants in age- and treatment-related CH, of which this paper presents a snapshot.

## Results

### Identifying somatic mutations in blood samples

We reasoned that low-coverage whole-genome sequencing of blood samples routinely carried out in cancer genomics projects may be repurposed to detect CH. Therefore, we obtained the DNA sequences of blood and tumor samples (paired samples) from two large cancer genomics cohorts. The first cohort comprised 3,785 paired samples obtained from metastatic solid cancer patients (metastasis cohort) sequenced at the whole-genome level^29^. The second included 8,530 paired samples collected from primary solid tumor patients (primary cohort) sequenced at the whole-exome level^28^.

Although possible, the identification of somatic mutations across the blood samples taken from the patients of these cohorts is extremely challenging due to the low coverage employed to sequence them. Therefore subclonal mutations are hard to distinguish from random sequencing errors. Moreover, somatic mutations at high variant allele frequencies and germline variants may also be confused if a somatic mutations calling is carried out on the blood sample alone. We reasoned that this problem could be overcome using the second (tumor) sample taken from the same patient as a reference of their germline genome. A comparison of the variants identified in the blood sample and the tumor sample with respect to the human reference genome would then reveal the somatic mutations specific to hematopoietic cells (Fig. 1a).

**Figure 1.**
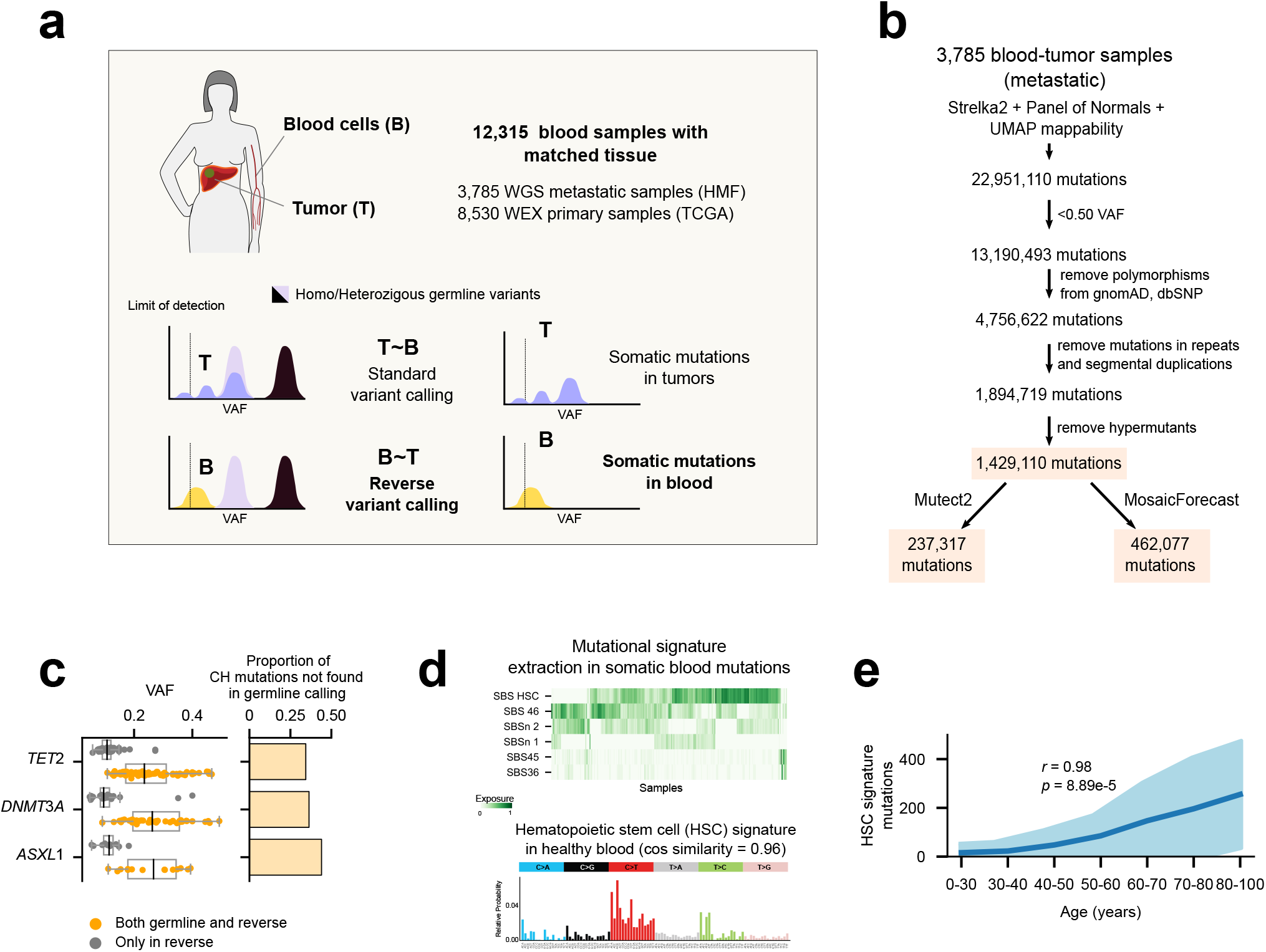
The reverse calling approach to detecting blood somatic mutations. (a) Conceptual depiction of the reverse calling approach. Somatic mutations in blood are identified by comparing variants in the blood/tumor paired samples taken from a cancer patient, with the tumor as the “germline” genome of the donor. Variants that are unique to hematopoietic cells and above the limit of detection of the bulk sequencing (i.e., shared by enough number of hematopoietic cells) will be identified by the approach. We applied this approach to two cohorts of primary and metastasis tumors totalling 12,315 blood donors with no known hematologic phenotype. (b) Flowchart of the reverse calling and filtering approach. The numbers correspond to the putative somatic mutations remaining in the dataset (full, mosaic or mutect) when the successive filtering steps are applied to variants detected in the metastasis dataset. (c) Comparison of the somatic mutations and germline variants identified by the reverse calling approach and a traditional one-sample variant calling across blood samples of donors in the metastasis cohort. Left-hand boxplots represent the distribution of variant allele frequency (VAF) of somatic mutations affecting three well-known CH driver genes, which are identified by both approaches. The distribution of VAF of somatic mutations identified through the application of the reverse calling starts at lower values than those detected through a traditional one-sample calling. Right-hand barplots illustrate the fraction of mutations affecting each gene that are identified only by the reverse calling approach. (d) Mutational signatures active in the blood samples of donors across the metastasis cohort (identified using the mosaic set). The top heatmap represents the activity of the 6 mutational signatures extracted across all samples. The bottom barplot represents the mutational profile of tri-nucleotide probabilities of one of the signatures extracted from the cohort which highly resembles (cosine similarity = 0.96) that of a signature active in healthy hematopoietic stem cells (HSCs). (e) Relationship between the number of mutations contributed by the HSC signature across blood samples in the metastasis cohort and the age of their donors. Donors are binned according to their age, with the mean activity of the signature across donors of each bin represented as the dark blue line, and its standard deviation represented in light blue color. A significant positive correlation between the two variables is apparent.

We thus --inspired by a previous approach to identify early mutations in the development of the hematopoietic system^30^-- implemented a pipeline to systematically carry out this “reverse” somatic mutation calling on the paired samples of the two cohorts (Fig. 1b; Fig. S1a; Supp. Note). First, blood mutations are identified using a somatic mutation caller widely employed in cancer genomics studies^31^, and a set of filters of the quality of the calls are applied to guarantee that these are true somatic mutations rather than germline variants or random sequencing errors. In the metastasis cohort, this yields ∼1 million candidate whole-genome somatic mutations across 3,785 patients. We call this the full catalog of somatic mutations. Two further filtered sets are obtained applying one of two criteria (Fig. 1b): mutations also identified by a second widely-employed somatic caller^32^ (mutect catalog), or mutations identified by MosaicForecast, an algorithm trained to detect likely somatic mutations using phased mutations^33^ (mosaic catalog; Fig. S1b). Importantly, the reverse calling approach empowers the detection of variants in known CH genes at values of variant allele frequency unattainable by a typical germline calling on a single whole-genome sequenced blood sample (Fig. 1c). Taking as example three well known CH drivers (TET2, DNMT3A and ASXL1), around 30% of all mutations identified by the reverse calling are missed by a germline calling.

In summary, the reverse calling approach identifies somatic mutation with higher sensitivity and specificity. On the one hand, mutations at lower VAF may be identified (higher sensitivity), as explained above. On the other, it reduces the number of germline variants falsely called as blood somatic mutations (higher specificity).

### Somatic mutations in blood samples show evidences of clonal hematopoiesis

Only variants shared by enough number of blood cells --those that derive from the clonal expansion underlying CH-- would appear above the limit of detection of the sequencing. Therefore, the detection of these somatic mutations through this reverse calling approach is a first evidence of CH in the samples of both cohorts (Fig. 1b and Fig. S1a).

We expect that these mutations exhibit a tri-nucleotide profile characteristic of variants spontaneously appearing as HSCs divide^34^. The identification of mutational signatures active in the blood samples of the metastatic cohort yielded 6 distinct profiles. Some of these are similar to signatures previously associated with sequencing artifacts^35^ (Fig. S1c,d; Table S1). Nevertheless, the most pervasive mutational signature in the cohort shows a profile that is virtually identical (cosine similarity = 0.96) to that of the known hematopoiesis signature (Fig. 1d). This constitutes further evidence that a set of somatic mutations contributed by hematopoiesis are present across these healthy blood samples. Moreover, it is further indication that clonal hematopoiesis has occurred across at least some of the donors.

We also expect that blood somatic mutations contributed by HSC divisions increase with the age of the donors^34,36^. First, the chance of appearance of a CH mutation (a mutation affecting a CH driver), and in consequence the chance of the expansion of a HSC clone, increases with age. Second, the number of hematopoietic mutations in this HSC clone founder (which become amplified due to the clonal expansion), also increases with age, because hematopoietic mutations are acquired at a steady rate with every HSC division. Third, the longer the time elapsed between the beginning of the clonal expansion and the obtention of the sample (which naturally increases with the donor’s age), the higher the VAF of the hematopoietic mutations, and the likelihood that they rise above the limit of detection of bulk sequencing. In agreement with this expectation, we observed that the number of hematopoiesis mutations identified in the metastasis cohort applying the reverse calling approach increases with the age of the donor (Fig. 1e; Fig. S1d illustrates the relationship for all phased mutations). On the contrary, the number of mutations contributed by the other signatures extracted from the cohort does not increase steadily with age (Fig. S1e).

In summary, several lines of evidence provide support to the reverse calling approach as an efficient method to identify somatic mutations in blood samples of patients with CH when a paired tissue sample is available.

### Discovery of clonal hematopoiesis drivers

We reasoned that, as is the case in the clonal expansion related to tumorigenesis^25,37^, the mutational patterns of CH-associated genes should exhibit signals of positive selection across donor blood samples. Therefore, methods that have been developed over the years to identify these signals of positive selection in cancer^25,37–40^ could be applied to somatic mutations in blood samples to identify the genes with significant deviations from their expected patterns of mutations. Anchored in these methods, cancer genomics researchers have set the goal of uncovering the compendium of cancer driver genes. Analogously, exploiting these methods empowers us to open a roadmap to the compendium of CH driver genes.

To test this concept, we applied the IntOGen pipeline^25^ (which runs seven state-of-the-art driver discovery methods^41–47^ and combines their results) to blood somatic mutations in the primary and metastatic cohorts (Fig. 2a). Each of these methods identifies one or more signals of positive selection (e.g., abnormally high recurrence of mutations, unexpected clustering of mutations in certain regions of the gene, or exceptionally high functional impact of the observed mutations) in the mutational pattern of genes across samples (Fig. 2a and Supp. Note). False positive genes identified by a particular method are filtered out by the combination of their outputs through a voting-based approach^25^. Finally, 15 genes that are significant according to the combination approach are filtered out as they are deemed suspicious after a careful vetting that considers gene expression across HSCs, somatic hypermutation processes, common sequencing artifacts and frequent false positive genes of the driver discovery process (Supp. Note).

**Figure 2.**
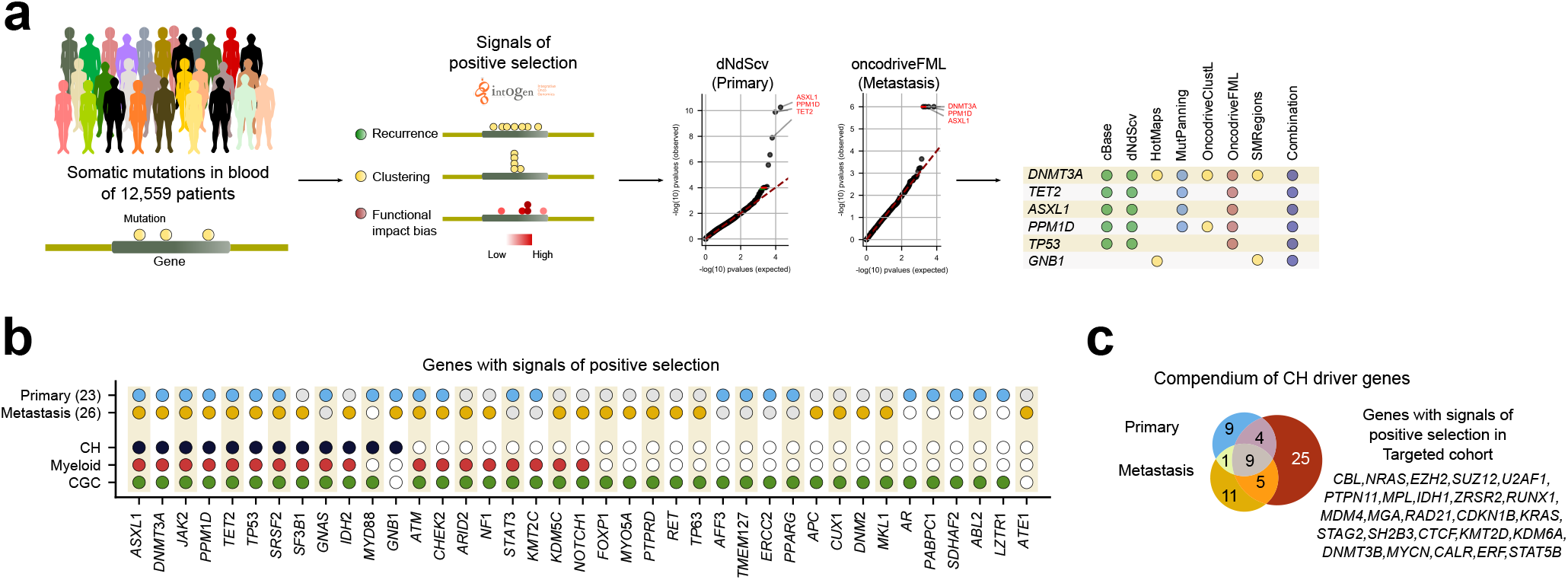
Discovery of clonal hematopoiesis driver genes. (a) Summary of the discovery analysis applied to blood somatic mutations detected across the primary and metastasis cohorts. The (differently filtered) sets of blood somatic mutations identified across all donors of a cohort were the input data for the analysis. Seven state-of-the-art driver discovery methods probing different signals of positive selection were applied (via the IntOGen pipeline) to each dataset of mutations. The IntOGen pipeline also handled the combination of the output of the seven methods to yield a unified list of CH driver genes in each cohort (details in Supp. Note). (b) CH driver genes discovered across the primary and metastasis cohorts. Genes known to be involved in CH, myeloid malignancies or tumorigenesis in general are labeled. (c) The compendium of CH driver genes is completed through the discovery carried out across blood samples in the targeted cohort (24,146 donors). The overlap in the lists of gene discovered across the three cohorts are shown in the figure. The discovery of CH drivers in this targeted cohort was carried out using a subset of the methods employed in the analysis of the primary and metastasis cohorts (see Supp. Note). The union of the lists of CH drivers discovered in these three cohorts (64 genes) integrate the CH drivers compendium presented in Figure S2, Table S2 and available through www.intogen.org/ch.

The lists of CH drivers are composed by 26 and 23 genes in the metastasis and the primary cohorts, respectively (Fig. 2b; Table S2). Employing a list including 15 well-known CH-related genes, identified across several studies as ground truth of clonal hematopoiesis drivers^48^ (CH known drivers), we found that most CH known drivers are detected in at least one of the two cohorts. Only CBL, NRAS, and KRAS appear below the statistical power of the IntOGen pipeline in these two cohorts. Nevertheless, all known CH drivers are identified when signals of positive selection are probed across 24,146 targeted-sequenced paired blood/tumor samples^17,49^ (targeted cohort) in which a mutation calling filtering variants in common with the tumor sample has been carried out (Fig. 2c; Supp. Note). Other genes with signals of positive selection, like ATM and CHEK2 are known to drive myeloid malignancies^17^. A third group of genes, such as CUX1 (mentioned in several studies as possibly related to CH)^13,21^, ABL2, FOXP1 or TP63 while known cancer drivers^50^, have not been previously proved to be involved in CH. These results, thus link these known cancer genes to CH. While some genes may be under the limit of detection of the methods of driver discovery in one of the two cohorts, ten genes exhibit signals of positive selection in both cohorts and also in the targeted cohort (Fig. 2c and Fig. S2).

Therefore, the identification of signals of positive selection in the pattern of somatic mutations of the genes across blood samples of healthy individuals is an effective way to discover CH-related genes, it recovers most known CH genes and has the power to discover new ones. This compendium --the snapshot presented in this work-- comprises the genes identified across the primary, the metastasis and the targeted cohorts and is available in Figure S2, Table S2 and through https://www.intogen.org/ch

### The drivers of clonal hematopoiesis

To further characterize discovered CH-related genes, we probed the association of their mutations with several physiological and clinical variables relevant to the development of CH (Fig. 3a,b). As previously reported^17^, across patients in the metastasis cohort, we found that the emergence of CH is positively influenced by age and by the exposure to cytotoxic (but not non-cytotoxic) treatments (Fig. 3a). We also recapitulated the prior knowledge that mutations in certain genes, such as PPM1D and CHEK2 are positively associated with the exposure to platinum-based drugs (Fig. 3b). Mutations in a group of three DNA-damage response genes (TP53, PPM1D, CHEK2) appear significantly associated with the exposure to platinum (Fig. 3b), probably because HSCs carrying them possess better survival rate than others when exposed to these DNA-damaging chemotherapeutics^2^. Interestingly, mutations in other genes involved in DNA-damage response, such as ATM do not appear to be positively associated with the exposure to platinum-based drugs.

**Figure 3.**
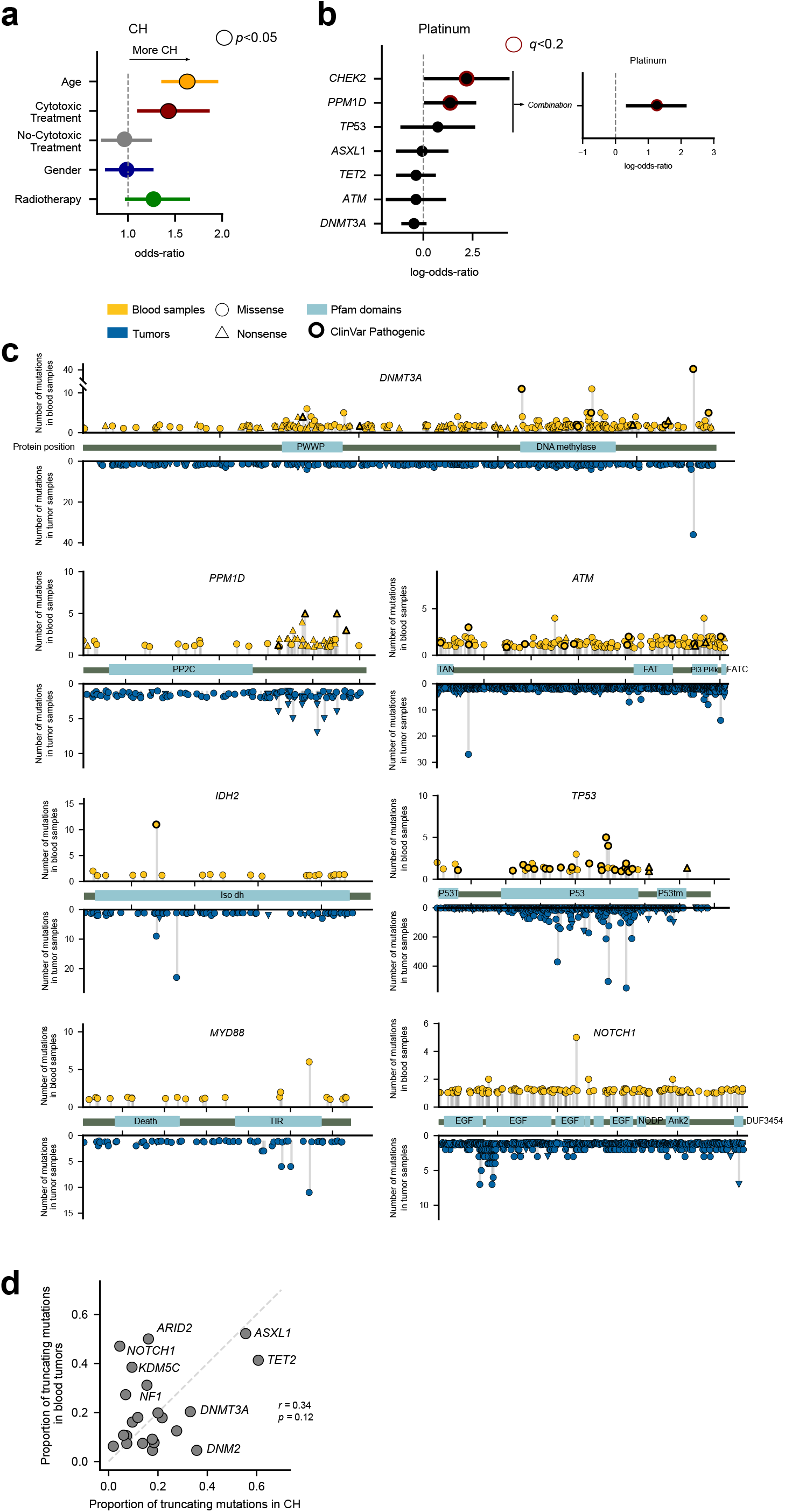
The drivers of clonal hematopoiesis. (a) Logistic regression showing the relationship between several factors and the development of CH across donors in the primary and metastasis cohorts. For this analysis, a donor is considered to suffer CH if they bear a non-silent mutation in a CH gene discovered in the analysis of the primary and/or metastasis cohorts here. The age of the donors in these cohorts as well as their prior exposure to cytotoxic therapies significantly increase their likelihood of presenting clonal hematopoiesis. (b) Logistic regression showing the relationship between the presence of mutations in several genes and the prior exposure of donors in the metastasis cohort to platinum-based therapies. Mutations in CHEK2 and PPM1D are significantly more likely across platinum-based exposed donors. (c) Distribution of blood somatic mutations affecting seven selected genes in the CH drivers compendium across donors of the primary and metastasis cohorts (above the horizontal axis) in comparison to those observed in the same genes across 28076 tumors^25^ (below the horizontal axis). (d) Relationship between the fraction of truncating variants identified in genes with 10 or more mutations across blood samples in the primary and metastasis cohorts and across several cohorts of tumors^25^.

We then asked whether the pattern of CH-related mutations of known cancer genes differ from that of their oncogenic mutations (Fig. 3c and Fig. S3). In the case of DNMT3A, one of the main hotspots of CH-related mutations (affecting residue 882) also appears recurrently mutated across tumors, while two other hotspots (residue 635 and 736) seem to be more specific of CH. In the case of TP53 mutations in both CH and cancer cases appear clustered across the DNA binding domain. The distribution of mutations of PPM1D is very similar across CH and cancer cases. In both scenarios, PPM1D truncating mutations close to the C-terminal yield a protein product lacking a degron, which is thus abnormally stable and results in the down-regulation of DNA-damage response and the proliferation of cells in the presence of such damage^51^. The distribution of mutations across CH and cancer cases is also very similar in the cases of MYD88 (with one dominant hotspot), but differ in IDH2. The pattern of mutations observed in these discovery CH genes across the primary and metastasis cohorts remarkably resemble those obtained across the targeted cohort (Fig. S4a). The distribution of mutations along the sequence of other genes in the compendium is shown in Figure S3.

While many CH drivers exhibit similar frequency of truncating mutations across both CH and myeloid cancer cases, in some, a clear enrichment (TET2, PPM1D) or depletion (NOTCH1, ARID2) of mutations with this consequence is observed across blood samples (Fig. 3d). Interestingly, the rate of truncating mutations in CH driver genes across donors of the primary and metastatic cohorts is very similar to that observed in the targeted cohort (Fig. S4b). The case of NOTCH1, mutations of which are related with the development of hematopoietic malignancies, such as ALL and CLL, could indicate that different constraints underlie the development of CH and these malignancies^52^. (Interestingly, the low share of truncating mutations of NOTCH1 is observed across the three cohorts analyzed; Fig. S4b.) The observed differences between CH and cancer may have their origin in the disparate array of mutational processes active in healthy blood and tumors, or in different evolutionary constraints.

### Detecting clonal hematopoiesis across ∼12,000 donors

Using the list of CH-related genes, we then identified all patients across the primary and metastatic cohorts with a potential protein-affecting somatic mutation in one of these genes (Fig. 4a). This way, we identified 1676 (19%) and 346 (9%) healthy blood donors who carried a potentially CH-related protein-affecting mutation in the primary and metastatic cohorts, respectively. The rate of mutations of the most frequently mutated CH genes varies between both cohorts (Fig. 4b), reflecting differences in mutational processes and evolutionary constraints related to CH emergence. The most frequent mutational hotspots affect JAK2 and DNMT3A (Fig. 4c). Interestingly, while more than three quarters of the patients across both cohorts present only one mutation affecting a CH gene, more than one are identified in 18% (Fig. 4d). These co-occurring mutations affect some CH-related genes more frequently (Fig. 4e; Fig. S5a). PPARG, SF3B1, PPM1D and TMEM127 are among them.

**Figure 4.**
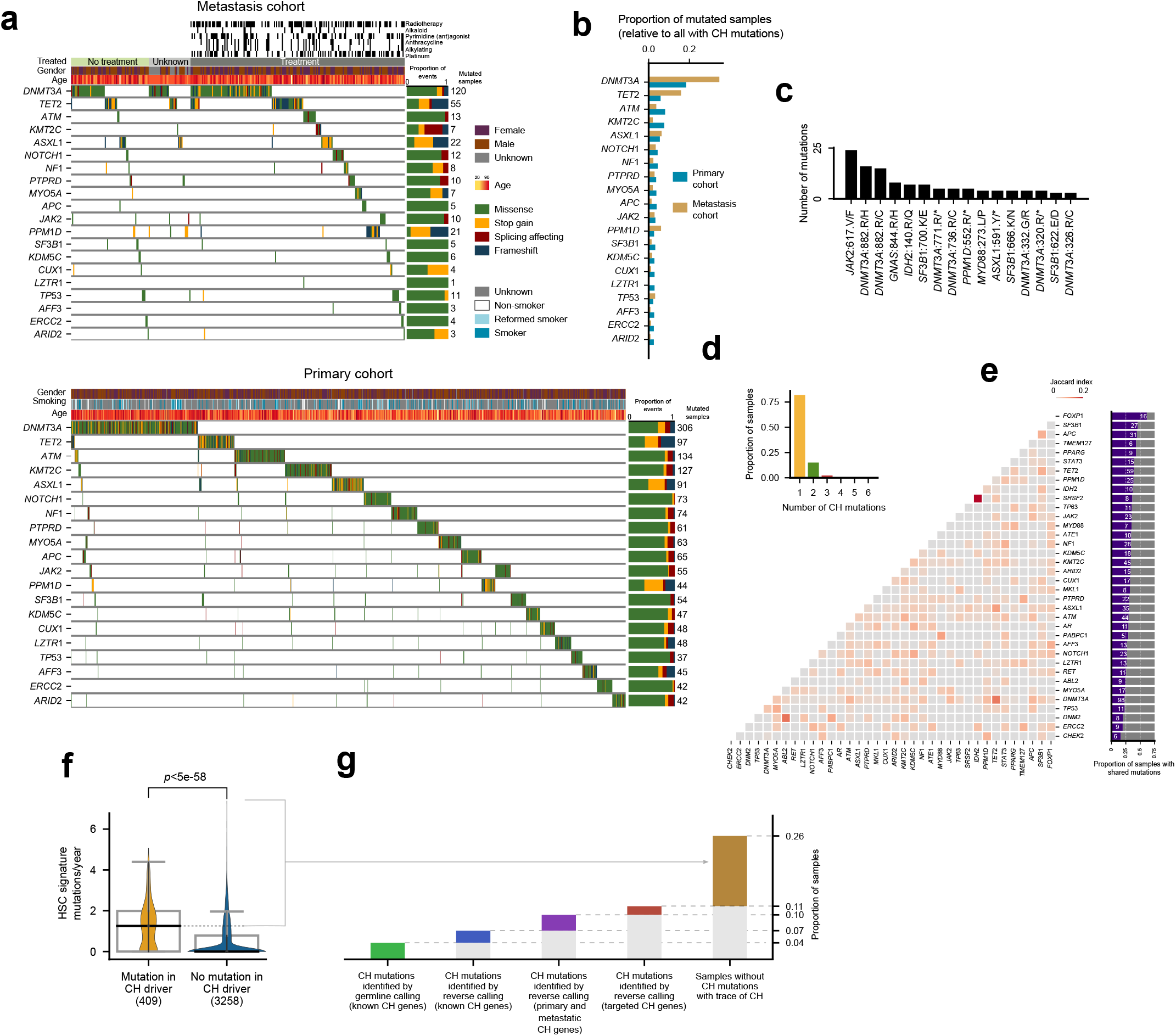
Clonal hematopoiesis across 12,000 donors. (a) Blood somatic mutations affecting the 20 more recurrently mutated genes in the CH drivers compendium across the metastasis (top heatmap) and primary (bottom heatmap) cohorts. Different types of mutations are represented with different colors and their observed frequency in each gene are summarized in the hand-right stacked barplots. The age, exposure to treatment and other clinically relevant features of the donors in each cohort are annotated above the heatmaps. The detected somatic mutations in these CH drivers define the presence of CH across these donors. (b) Frequency of mutation of CH drivers shown in (a) across the metastasis and primary cohorts. The frequency (number of donors with blood somatic mutations affecting the gene) is normalized by the total number of donors with mutations in genes in the CH drivers compendium. (c) The 16 most recurrently mutated hotspots in genes in the CH drivers compendium. (d) Number of donors in the two cohorts with different number of mutations in genes in the CH drivers compendium. (e) Frequency of co-occurring mutations in genes in the CH drivers compendium. The left-hand triangular matrix presents the co-occurrence of mutations between specific pairs of genes relative (Jaccard’s index) to their total mutational frequency. The right-hand barplot presents the frequency of each gene’s co-mutation with any other gene in the CH drivers compendium. (f) Distribution of the rate of hematopoietic mosaic mutations per year identified in donors of the metastasis cohort across (left) donors bearing a mutation in a gene identified as CH driver in the primary or metastasis cohorts and (right) donors with no detected mutations in any of these genes. The horizontal dashed line extends out of the median of the distribution of rate of mutations per year of age of the donors with mutations in at least one CH gene, representing the donors in the second group that are considered to be cases of clonal hematopoiesis (see next panel). The two distributions were compared using the two-tailed Wilcoxon-Mann-Whitney test. (g) Number of donors in the metastasis cohort with clonal hematopoiesis recognizable through different criteria. The first bar presents the number of donors identified if only mutations (detected in a one-sample calling) affecting a known CH gene are considered. The second bar counts the number of donors added if all variants identified through the reverse calling that affect known CH genes are included. The third bar comprises the donors added if mutations (identified through the reverse calling) affecting CH genes discovered across the primary or metastasis cohorts are taken into account. The fourth bar presents the number of donors added by considering also mutations affecting CH genes that are only discovered in the targeted cohort. Finally, the fifth bar contains the number of donors in the metastasis cohort with no mutation in any gene within the compendium of CH drivers, but with more hematopoiesis mutations per year of age of the donor than the median rate of hematopoiesis mutations across donors in the four previous groups.

The range of variant allele frequency (VAF) of the mutations in these genes reveals a wide spectrum of clonal expansion across CH cases (Fig. S5b). The rate of hematopoiesis mutations per year of age detected in patients with an identified mutation in a CH-related gene is significantly greater than that detected across patients without an identifiable CH-related mutation (Fig. 4f; Fig. S5c). The explanation for this finding is that hematopoietic mutations are more likely to appear above the threshold of detection of bulk sequencing the greater the CH clone. In samples carrying a mutation in a *bona fide* CH driver, it is more likely that this clone has expanded enough to identify a set of hitchhiking mutations through the reverse calling. Conversely, among samples without a CH-related mutation it is more likely that the clone is smaller or not present at all (detected hematopoietic mutations may be false positives of the reverse calling). In either case, the number of identified mutations is expected to be smaller across these patients.

We then set out to detect all CH cases across the metastasis (Fig. 4g) and primary cohorts (Fig. S5b). First, we determined that 141 CH cases in the metastasis cohort would be detected just by identifying somatic variants affecting genes in the list of known CH drivers on the bases of the blood germline calling (4% of the patients in the metastasis cohort). Using the reverse calling to identify somatic variants affecting these genes would add 99 CH cases (ascending to 7% of the total number). The addition of all CH-related genes to the compendium in this paper identifies 110 further CH cases (up to 10%), with 59 (ascending to 11%) more added if the set of CH driver genes identified across the targeted cohort is also considered. Across donors in the primary cohort, 25% are detected as CH cases following the same criteria (Fig. S5d).

Finally, we assumed that any sample with a rate of hematopoiesis mutations per year above the median of the distribution of values observed for samples carrying CH-related mutations is a case of CH, even in the absence of identified driver mutation (Fig. 4f). Thus, 562 (totalling 15%) blood samples in the metastasis cohort with no detectable CH-related mutations exhibit a rate of hematopoietic mutations comparable to that of samples with a mutation in a *bona fide* CH driver (Fig. 4g). We reasoned that at least some of these CH cases --with an appreciable clonal expansion-- could be driven by mutations affecting yet unidentified CH drivers or may have resulted from expansion of HSCs due to non-genetic mechanisms.

Still, some CH cases may be driven by non-coding mutations. Whole-genome sequenced blood samples may be employed to identify such non-coding driver mutations. This is not an easy task, as demonstrated by the search for non-coding cancer driver events^53,54^. The possibility is nevertheless opened by the reverse calling demonstrated here to set out to identify signals of positive selection in the observed pattern of mutations of different non-coding genomic elements. This is demonstrated with the results of OncodriveFML^41^, MutSigCV_NC^54^ and DriverPower^55^ on mosaic mutations in non-coding genomic elements (Fig. S6a and Table S3). The results of such analyses need to undergo a rigorous vetting process, as the distribution of mutations under neutrality in non-coding regions is still very difficult to model. Alternatively, the functional effect of mutations overlapping particular non-coding regulatory elements, such as the binding site of a transcription factor in an enhancer element, may be assessed. For example, Figure S6b illustrates the potential disruption of a binding site for RARA in an enhancer element potentially regulating TET2. Figure S6c (see more examples in Table S4) presents the potential creation of a SALL4 binding site in an enhancer regulating the expression of GNAS.

In summary, we show how the compendium of CH driver genes improves the identification of clonal hematopoiesis across large cohorts of donors, as a first step towards the study of all the molecular mechanisms behind this phenomenon.

## Discussion

The extent of CH across patients with no known hematologic phenotype is currently not well gauged, although population studies have revealed that it is probably higher than anticipated a few years ago^2,3,13,14,16,21,22^. Understanding this extent and comprehensively identifying CH across healthy donors is key to predicting potential future health hazards. One stepping stone in this path is the identification of all genes with mutations capable of driving CH. Moreover, the identification of all CH-related genes is a requisite to understand the mechanisms behind this process and its relationship with disease conditions, as has been done for mutations affecting chromatin remodelling and DNA damage response genes classically associated with the condition^2,16,17,51^. In this regard, the discovery of CH-related genes across populations of various ethnicities and with different lifestyles, will allow us to understand the different constraints faced by hematopoietic cells in their evolution and the outcome of each of them.

In this study, we have demonstrated that the existence of a second non-blood sample of the same donor refines the identification of somatic mutations in a blood sample, even though this is sequenced at low depth. The reverse calling implemented and tested here identifies blood somatic mutations with more sensitivity (across all discovery CH drivers) and more specificity (owing to the tumor paired sample) than a regular germline calling on a single blood sample. Importantly, the identification of mutational signatures active in a blood sample that may be the result of sequencing artifacts calls to caution when interpreting these blood mutations. For instance, a simple count of the number of somatic mutations identified should not be considered a signal of CH, as in some samples these may be artifactual mutations. Instead, we propose to use the mutations most likely contributed by the HSC mutational signature (Figure 4f,g).

We have also demonstrated that CH-related genes may be systematically and unbiasedly identified through the repurposing of tools aimed at identifying genes under positive selection in tumorigenesis. If these two elements --newly identified CH-related genes and the reverse mutation calling-- are combined, the identification of CH across donors gains in sensitivity. While the size of the cohorts employed here limit the power of the discovery of new CH drivers, and the mechanistic inferences that can be made from them, we envision that the application of this rationale to large tumor sequencing cohorts contribute to expand the list of CH drivers. This effort would benefit --as is apparent from the previous paragraph-- from deeper sequencing of the reference blood samples in cancer genomics studies. Moreover, the evidence that CH may be present in a substantial number of samples in the absence of mutations of genes in the compendium underlines the pressing need to extend the discovery of CH drivers. In this regard, an analysis that repurposes many more tumor/blood paired samples obtained in the context of cancer genomics projects following the approach demonstrated in this paper is of paramount importance.

One clear benefit of a compendium produced via a systematic driver discovery effort with respect to the identification of recurrently mutated suspicious genes is that it will consider only those with clear signals of positive selection. Therefore, mutated genes that are passengers to the CH process will not be considered, even if they are known to be involved in tumorigenesis in solid tissues (see examples in Supp. Note). This, in turn will result in a more accurate identification of CH cases across donors.

Although a set of CH-genes common to both cohorts is apparent from the discovery, a plethora of genes specific to each of them also appears. This is probably due to differences in both cohorts: primary vs metastatic tumors, many of which have been exposed to chemotherapies. Mutations in some CH-related genes are indeed known to provide an advantage to hematopoietic cells under exposure to certain cytotoxic treatments. Other differences, such as the different composition of both cohorts, in terms of human populations and tumor types represented may also have a bearing in the differences in CH-related genes discovered in each^56^. Further studies are needed to clarify this point, which the availability of the discovery presented here now make possible to undertake.

The unbiased snapshot of the compendium of CH drivers identified has a series of implications for both CH and cancer research. It may be directly employed in the research of the molecular mechanisms underlying CH in different scenarios. The list of 64 genes discovered can also be employed to refine the identification of the condition across human donors. Such donor-wise identification of CH would require the analysis of a single blood sample, identifying variants affecting the genes in the compendium. An important warning arising from this work is that not all blood mutations affecting cancer driver genes play a role in CH. Thus, the results from sequencing panels that include genes without signals of positive selection in CH need to be carefully interpreted. In the cancer research field, our results support the idea that sequencing cell-free DNA isolated from blood samples with the aim of identifying tumor mutations in circulating genetic material may produce false-positive results caused by the detection of CH mutations^57,58^.

Whereas the compendium of CH drivers is a key first step in the detection of CH across individuals, a second necessary step consists in evaluating the capability of individual mutations in CH drivers to provide a selective advantage to HSCs. If only mutations with experimentally validated effect on CH or identified through epidemiological studies are considered as CH drivers, the prevalence of CH is underestimated. PPM1D, whose C-terminal truncating mutations are currently considered as CH drivers, provides a good example. It is possible that missense mutations affecting the degron that is lost in the truncation are also drivers of CH. On the other hand, taking into consideration all mutations affecting CH drivers probably leads to an overestimation of CH. Again, approaches developed in cancer genomics with the aim of identifying driver mutations may become useful in this task^59,60^.

In summary, the results shown here open up the possibility to repurpose the tens of thousands of blood and paired tissue samples collected across hundreds of cancer patients cohorts to expand the discovery of CH-related genes and to improve the identification of CH across their donors.

## Methods

### Sequences of samples from the primary and metastasis cohorts

The sequences of solid tumors and their paired blood samples (BAM files) were obtained from the Genomic Data Commons (GDC^61^) portal for the primary cohort (N=8,530) and from the Hartwig Medical Foundation (HMF^29^) repository for the metastatic cohort (N=3,785).

### HMF gemline calling

The germline variant calls of the HaplotypeCaller^62^ for the metastatic cohort were obtained from HMF^29^. All mutations, independently of the quality filters, were used to compare the sensitivity of this germline calling with the reverse calling developed in the paper (see below). This produces very conservative estimations.

### Detecting somatic mutations in blood samples across the primary and metastasis cohort (reverse calling)

The variant calling was carried out using the Google Cloud Platform (metastasis cohort) and in our in-house computer cluster (primary cohort). Briefly, the matched blood and tumoral BAM files --masked and deduplicated using GATK^62^-- of 3,785 whole-genome patients (metastasis cohort) and 8,530 whole-exome sequenced samples (primary cohort) were obtained as described above. The variant calling was carried using Strelka2^31^ with the blood sample as the tumoral input and the tumor sample as control (reverse calling). In the case of patients with more than one tumor sample, one of them was selected and included in the calling. All variants with two or more supporting reads matching the caller PASS filter and with VAF<0.5 were kept. Mutations in lowly mappable regions as defined by the DUST algorithm^63^ (k=30) and UMAP^64^ (36-kmers) were excluded. Contiguous variants were merged into double-base substitutions. Variants with greater frequency across the cohorts than the DNMT3A R882H or JAKV617E hotspot in a cohort-specific Panel of Normals (obtained from GDC and HMF for the primary and metastatic) and in gnomAD^65^ v2.1 were removed. This was equivalent to discard variants present in these datasets with an allele frequency greater than 0.002 in PoN TCGA, 0.008 in PoN HMF and 0.0003 in gnomAD v2.1. Additionally, common SNPs as defined by the snp151Common UCSC track^66^ and dbSNP^67^ were excluded. Mutations within segmental duplications, simple repeats and masked regions as defined in UCSC tracks were also removed. Finally, samples with the mutation count in the 97.5 percentile of the mutation burden across the cohort were deemed unreliable and excluded for further analyses. We call the set of variants obtained after the application of these filters the **full set**.

Two more conservative subsets were generated from the full set in the primary and metastasis cohorts. The first (**mutect set**) comprises only variants that were also called by Mutect2^32^ (only for the metastatic cohort). Second, we applied MosaicForecast^33^, a software designed to phase mutations to polymorphisms with the aim of identifying somatic mutations in a small number of cells and also predict mosaicism for the unphased ones with a random forest classifier. As a result, we obtained a subset of mosaic-phased mutations, and a subset mutations likely to be somatic (**mosaic set**). In the primary cohort, only the mosaic set was obtained through filtering of the full set.

### Blood somatic mutations in targeted-sequenced samples

Somatic blood mutations identified across 24,146 targeted-sequenced blood samples^17^ were directly obtained from cBioportal^68^.

### Detection of mutational signatures

To identify mutational signatures active in the metastasis cohort, we employed the mosaic set and applied a non-negative matrix factorization approach^69^, using the SigProfilerJulia (bitbucket.org/bbglab/sigprofilerjulia) implementation done in our lab^70^ of the algorithm developed by Alexandrov *et al*. (2013)^71^. Only samples with more than 100 mutations were included in the analysis. The resulting signatures were then compared to the PCAWG COSMIC V3^35^ catalog using the cosine similarity measure. No signature was extracted from the mutations identified in the primary (exome-sequenced) cohort due to their low numbers.

Whole-genome somatic variants of 23 blood samples from healthy donors of different ages were obtained from Osorio *et al*. (2018)^34^. The Hematopoietic Stem Cell Signature (HSC signature^34^) was computed as the average number of mutations observed across the 23 healthy blood samples in each of the 96 tri-nucleotide channels normalized by the total number of mutations observed.

### Discovering the compendium of CH driver genes

The discovery of genes with signals of positive selection was carried out using the IntOGen pipeline^25^. Briefly, the IntOGen pipeline implements seven complementary methods to identify signals of positive selection in the mutational pattern of genes and integrates their outputs. The IntOGen pipeline first pre-processes the somatic mutations across samples to filter out hypermutators, map all mutations to the GRCh38 assembly of the human genome and retrieve information necessary for the operation of the seven driver detection methods. Then, the methods are executed and their outputs combined using a weighted voting approach with weights adjusted depending on the credibility awarded to each method. Finally, in a post-processing step, spurious genes that result from known artifacts are automatically filtered out (see Supp. Note). The version of the pipeline used in this study is described at length at www.intogen.org/faq and in *Martinez-Jimenez et al*. (2020)^25^.

The IntOGen pipeline was run on the full set, the mutect set (metastatic cohort) and the mosaic set of mutations independently. Subsequently, genes that were identified as having signals of positive selection only in the full set were required to possess extra evidence (either identified by the pipeline run on a filtered set, or included within the Cancer Gene Census^50^) to be included in the final list. To compare CH-related genes according to this unbiased discovery to the prior knowledge on the genetics of this process, we used i) a list of genes involved in CH (ground truth of known CH genes^48^), ii) genes known to drive myeloid malignancies^17,21^, and iii) all genes annotated in the Cancer Gene Census^50^.

Only a subset of the methods (capable of building a background mutations model from the segment of the exome probed in the panel) were run on the set of somatic mutations identified in the blood samples of the targeted cohort. OncodriveCLUSTL, OncodriveFML, DnDscv (without genome-wide mutation rate covariates, as in ref^72^), and HotMaps were run through the IntOGen pipeline, and their individual outputs collected. Genes significant (with a FDR cutoff of 0.01) in the analysis of any method (that is a union of the lists) were considered CH drivers in this cohort.

The final snapshot of the compendium of CH driver genes was integrated by the union of the lists of genes identified across the three cohorts.

### Identification of blood samples with clonal hematopoiesis

To identify individual donors in the metastasis cohort with clonal hematopoiesis, we considered all mutations that putatively affected the protein sequence of any gene discovered as CH-related across the cohort in the present study (separated in the different categories presented in Fig. 4g). We then computed the median rate of hematopoiesis mutations per year of age across the blood samples of these donors. All donors with no mutation in a discovery CH gene but with a rate of hematopoiesis mutations per year of age greater than this median value were also considered to bear CH (the final group in Fig. 4g).

### Logistic regressions

Similar to a previous work^17^, we used multivariable logistic regression to assess the association between clonal hematopoiesis and therapy, age and gender. We also used it to compute the association between mutations in specific genes (or groups thereof) and the exposure of donors to specific chemotherapeutic drugs. Multiple test correction (Benjamini-Hochberg FDR) was used for gene-specific analyses.

### Identifying expressed CH-related genes

We computed the distribution of the expression of each gene across bone marrow CD34+ cells (GSE96811^73^). These cells are phenotypically the closest to the HSCs. We deemed a gene expressed across the cells when the maximum value of its distribution was above 15 fpkm.

### Comparison of blood somatic mutations with tumor mutations

The distribution of mutations in CH driver genes observed across blood samples from the primary and metastasis cohorts was compared to that observed across hematopoietic malignancies in IntOGen^25^. ClinVar pathogenic and likely pathogenic variants were obtained from ref^74^.

### Non-coding blood somatic variants in CH

Three state-of-the-art methods designed to detect positive selection in the mutational patterns of non-coding genomic elements (OncodriveFML^41^, DriverPower^55^, MutSigCV_NC^53^) were run with default parameters. The non-coding genomic elements were obtained from the PanCancer Analysis of Whole Genomes (PCAWG)^26^. A FDR cutoff of 0.2 was applied.

The set of transcription factor (TF) binding motifs was obtained from (ref^75^). Models with A,B and C qualities were kept. Only TF expressed in CD34+ cells according to GSE96811 were allowed in the analysis. H3K27ac ChIP data for CD34+ samples was obtained from ENCODE^76^. Enhancer element coordinates, as well as their defined target genes, were retrieved from geneHancer^77^ via the UCSC genome browser. Briefly, mutations intersecting with H3K27ac peaks and an enhancer defined by geneHancer were expanded 15bp upstream and downstream. Then, using FIMO^78^ the binding affinity of these sequences was determined for both the mutant and the reference allele. When the significance of the binding was less than 0.0001 in the reference but not in the mutant, we labeled the instance as disruption (and creation, if the case is the opposite). We retained only results for which the gene closest to the disruption/creation of a TF is a CH driver. Visualization of the genomic context of the mutations represented in Figure S6 was performed using pyGenomeTracks^79^.

### Code availability

The programs required for the variant calling are all open source, as is the IntOGen pipeline (available at www.intogen.org), and the programs used in the analysis of CH non-coding mutations (listed in the previous section). All other analyses described in the paper were implemented ad hoc in Python. The command lines employed in the calling and all code needed to reproduce all other analyses will be available in a public repository at the time of publication.

### Data availability

The data employed in the paper is available through different sources. Whole genome sequences of tumor and blood samples in the metastasis cohort are available from the Hartwig Medical Foundation for academic research upon request (https://www.hartwigmedicalfoundation.nl/en). Whole exome sequences of tumor and blood samples from the TCGA cohort are also available upon request through dbGAP or GDC. Panel-sequenced data from the IMPACT targeted cohort is available through cBioPortal. Other datasets employed in specific analyses are described in prior sections of these Methods.

## Contributions

O.P., A.G.-P. and N.L.-B. designed the project. O.P. carried out all the analyses and prepared the figures. O.P and I.R-S implemented the pipeline in GCP. I.R-S implemented the web site. N.L.-B. and A.G.-P drafted the manuscript. O.P., A.G.-P. and N.L.-B. edited the manuscript. A.G.-P. and N.L.-B. supervised the project.

## Acknowledgements

The authors wish to thank fruitful discussions of the results of the paper with Jose J. Fuster. N.L-B. acknowledges funding from the European Research Council (consolidator grant 682398) and ERDF/Spanish Ministry of Science, Innovation and Universities - Spanish State Research Agency/DamReMap Project (RTI2018-094095-B-I00) and Asociación Española Contra el Cáncer (AECC) (GC16173697BIGA). IRB Barcelona is a recipient of a Severo Ochoa Centre of Excellence Award from the Spanish Ministry of Economy and Competitiveness (MINECO; Government of Spain) and is supported by CERCA (Generalitat de Catalunya). O.P. is the recipient of a BIST PhD fellowship supported by the Secretariat for Universities and Research of the Ministry of Business and Knowledge of the Government of Catalonia, and the Barcelona Institute of Science and Technology (BIST). This publication and the underlying research are partly facilitated by Hartwig Medical Foundation and the Center for Personalized Cancer Treatment (CPCT) which have generated, analysed and made available data for this research. We would like to thank Paul Wolfe from Hartwig Medical Foundation for his guidance in Google Cloud Platform usage. This publication and the underlying research are also partly facilitated by data collected and made public by The Cancer Genome Atlas network.

---

**Figure S1.**
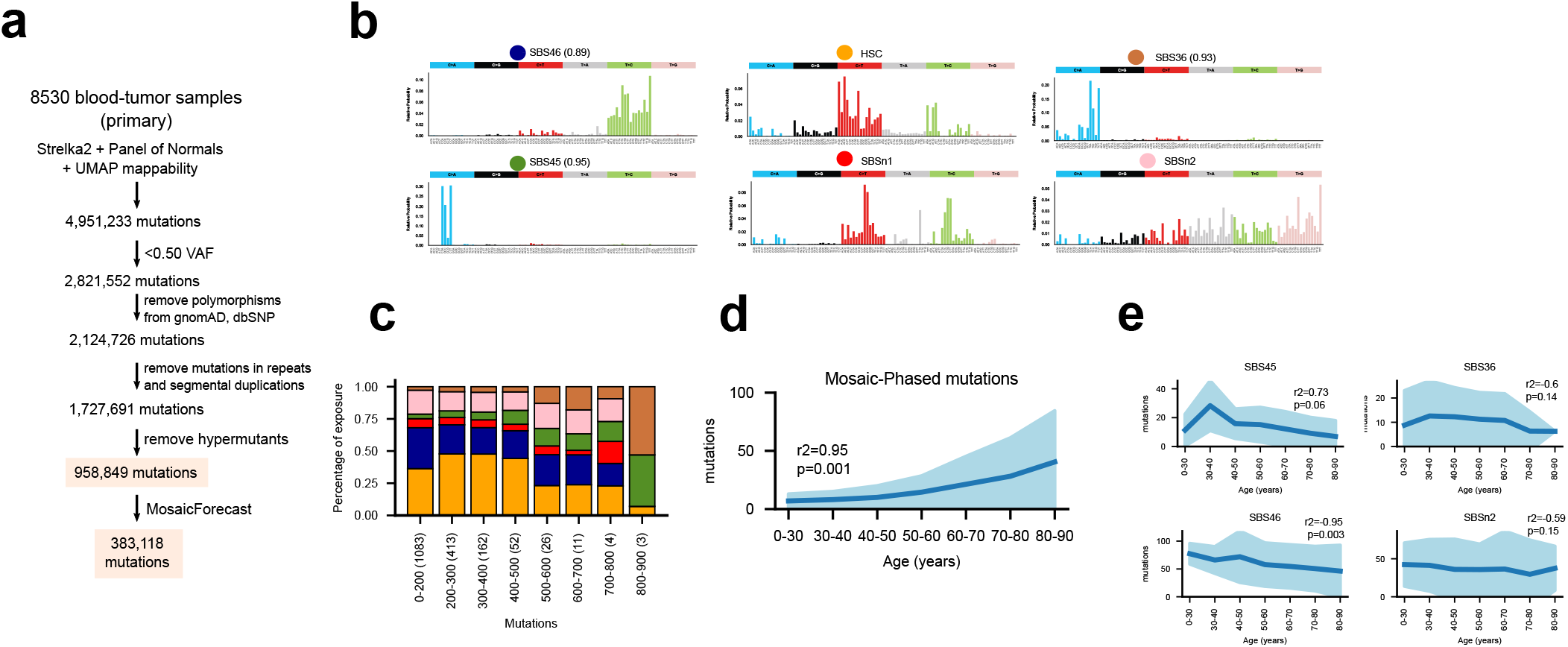
Identification of blood somatic mutations across the primary cohort. (a) Flowchart of the reverse calling and filtering approach in its application to the primary cohort. In this case only one filter of the set of mutations is applied. Therefore, two sets of somatic mutations are derived from the reverse calling pipeline: the full set and the mosaic set. (b) Tri-nucleotide profiles of mutational signatures extracted from the somatic mutations in the metastatic cohort. (c) Distribution of the activity of different mutational signatures active in the metastasis cohort across samples with different burden of somatic mutations. (d) Significant positive correlation between the number of phased mutations (yielded by the MosaicForecast algorithm) and the age of donors in the metastasis cohort. (The same general trend is shown in Fig. 1e for HSC signature mutations.) (e) Lack of significant positive correlation between the number of mutations contributed by different mutational signatures (except the corresponding to the HSC signature, depicted in Fig. 1e) and the age of donors in the metastasis cohort.

**Figure S2.**
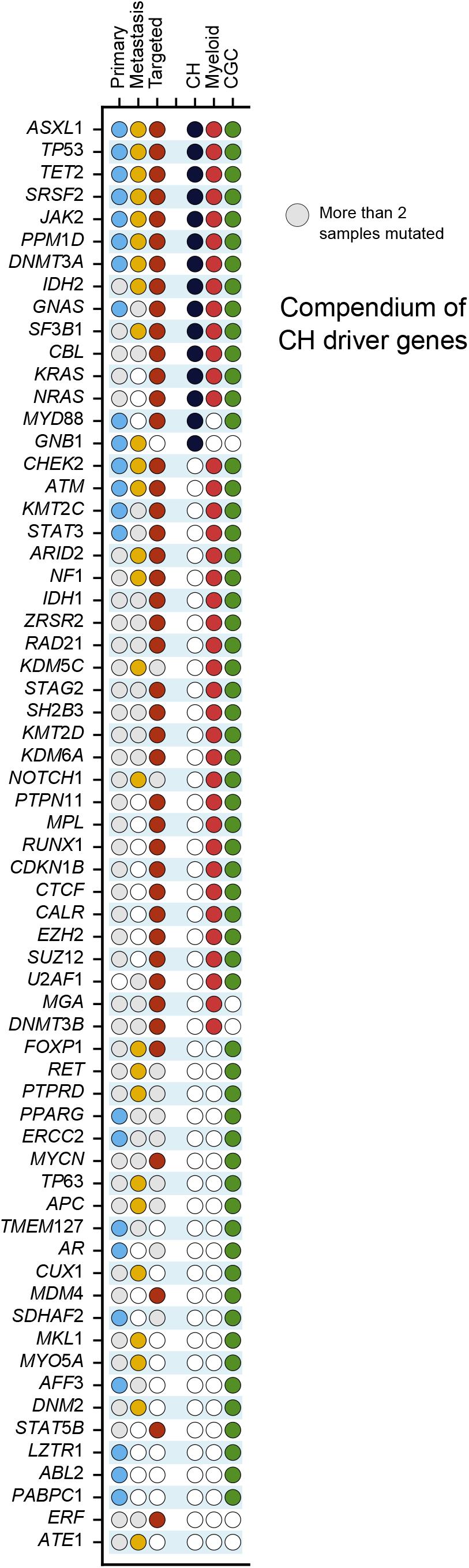
The compendium of CH driver genes. The compendium of CH driver genes presented as an annotated list of genes similar to that in Fig. 2b.

**Figure S3.**
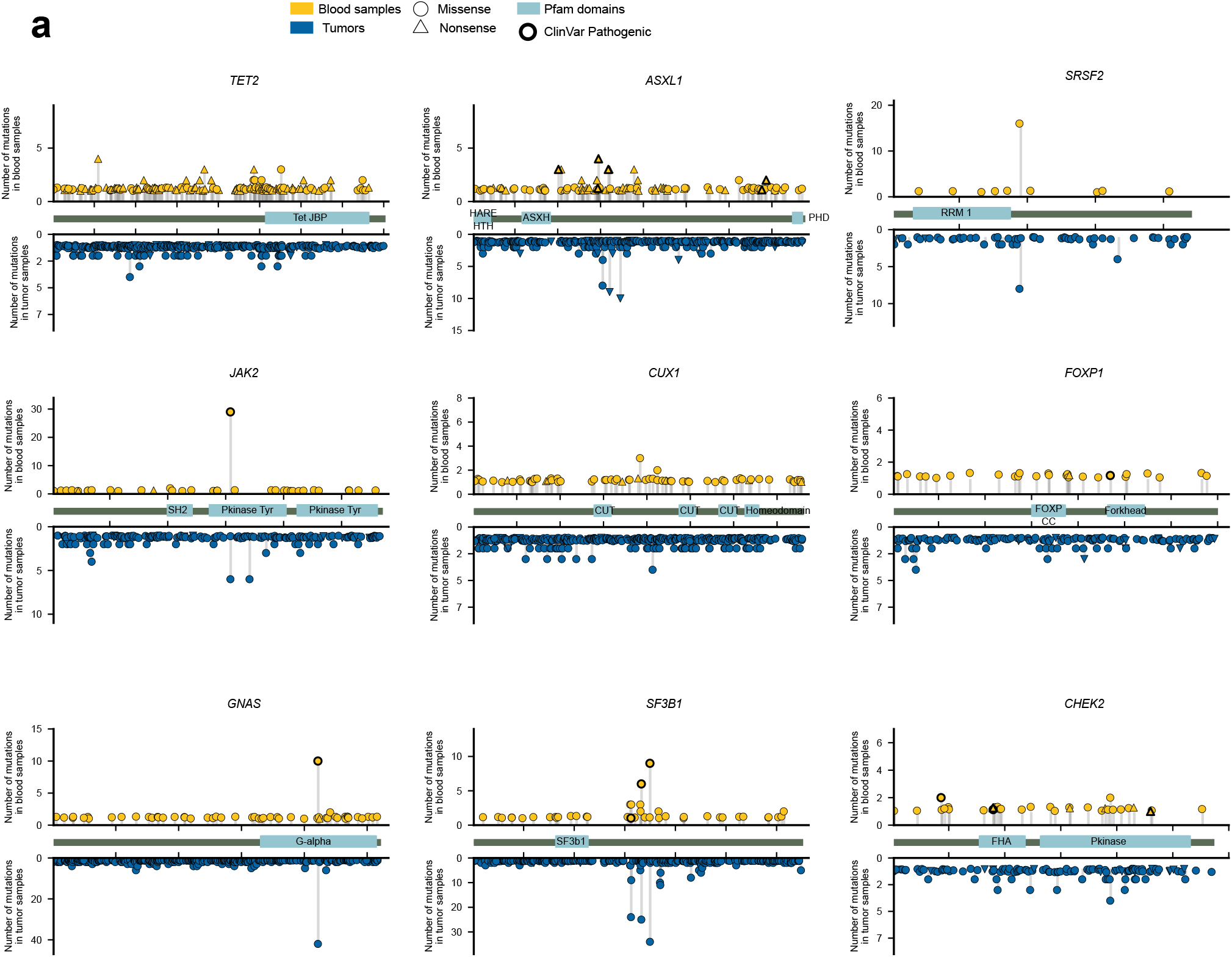
Distribution of blood and tumor mutations in CH driver genes. (c) Distribution of blood somatic mutations affecting seven selected genes in the CH drivers compendium (in addition to those presented in Fig. 3b) across donors of the primary and metastasis cohorts (above the horizontal axis) in comparison to those observed in the same genes across 28076 tumors^25^ (below the horizontal axis).

**Figure S4.**
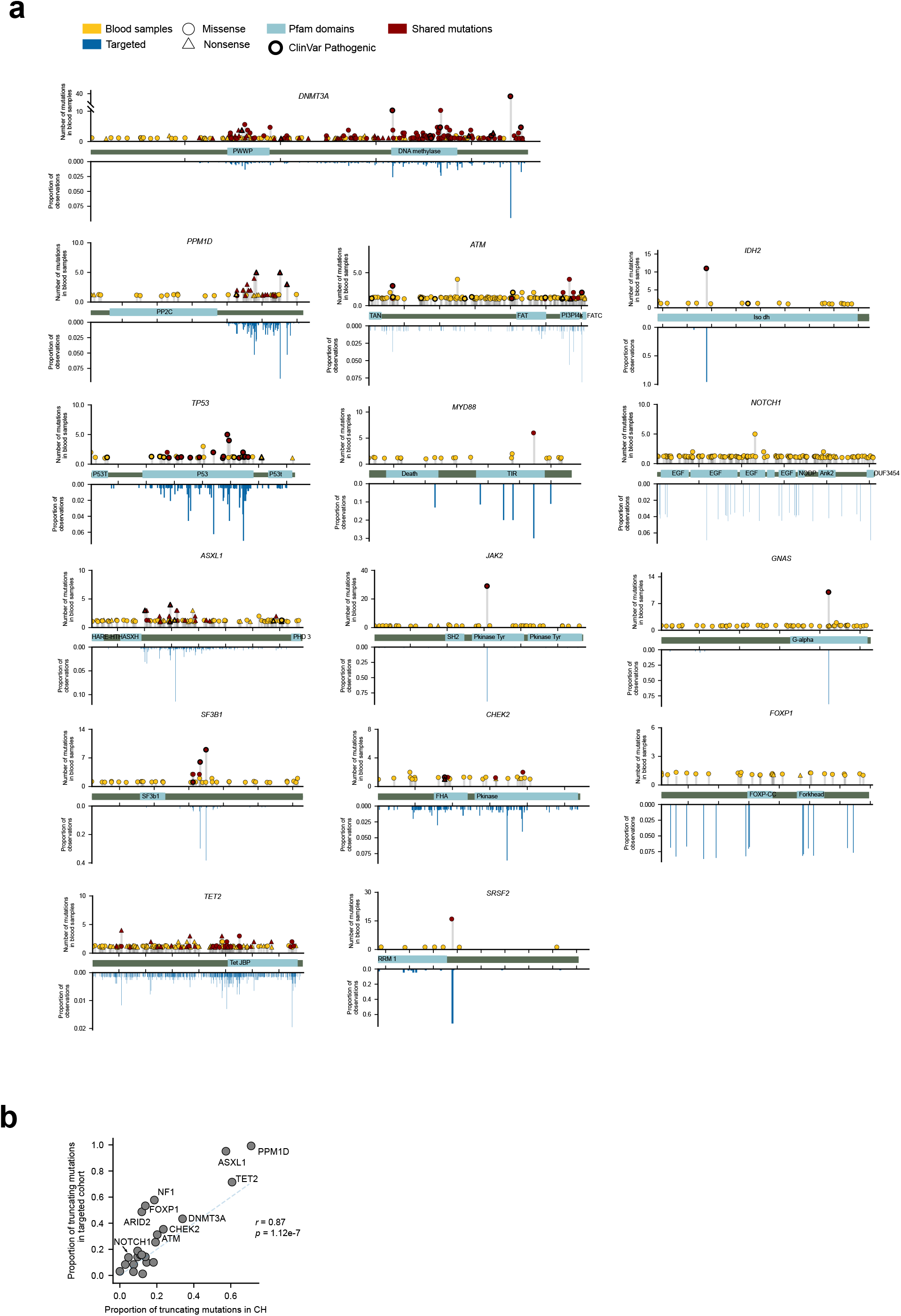
Distribution of blood mutations in CH driver genes across different cohorts. (a) Distribution of blood somatic mutations along the sequence of 15 CH driver genes (presented in Fig. 3b and Fig. S3) across the primary and metastasis cohorts (above the x-axis) compared to those identified across the targeted cohort (below the x-axis). Proportions with respect to the total number of mutations are shown across the targeted cohort, while the absolute numbers are illustrated for primary and metastasis cohorts. (b) Fraction of truncating mutations in CH driver genes with 10 or more mutations across the primary and metastatic (x-axis) and the targeted (y-axis) cohorts.

**Figure S5.**
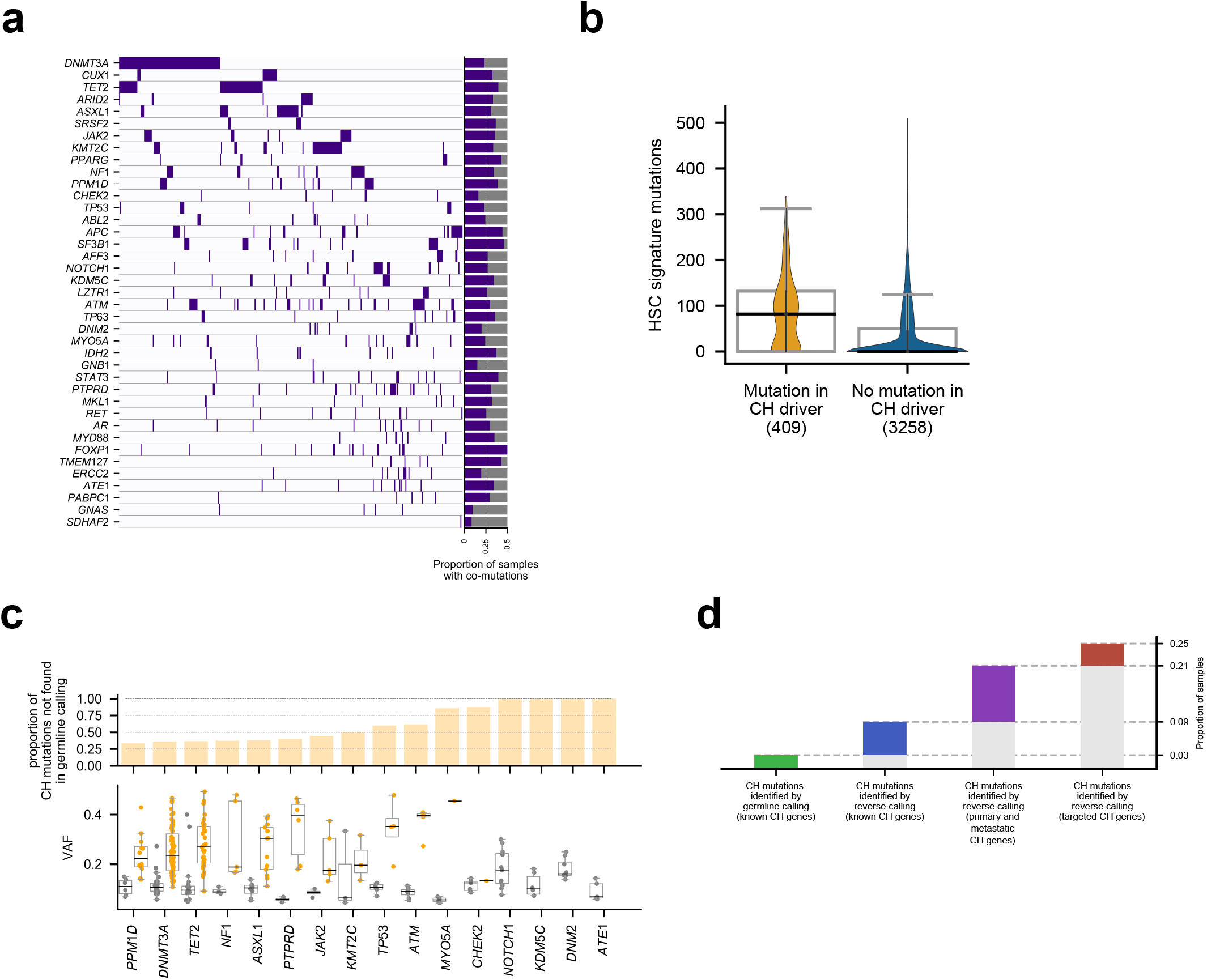
Detection of clonal hematopoiesis across donors. (a) Mutation co-occurrence across pairs of CH driver genes. (b) Distribution of number of hematopoiesis mutations of donors with mutations in a CH driver gene (identified across the primary or metastasis cohorts) and those with no mutation in CH driver genes. (c) Comparison of the somatic mutations and germline variants identified by the reverse calling approach and a traditional one-sample variant calling across blood samples of donors in the metastasis cohort. Analogous to Figure 1c, but including more known and discovered CH driver genes. (d) Number of donors in the primary cohort with clonal hematopoiesis recognizable through different criteria. Similar to Figure 4g, but only with the first four sets of donors. The fifth set is not available due to the extreme difficulty of extracting a mutational signature with the low numbers provided by whole-exome sequencing.

**Figure S6.**
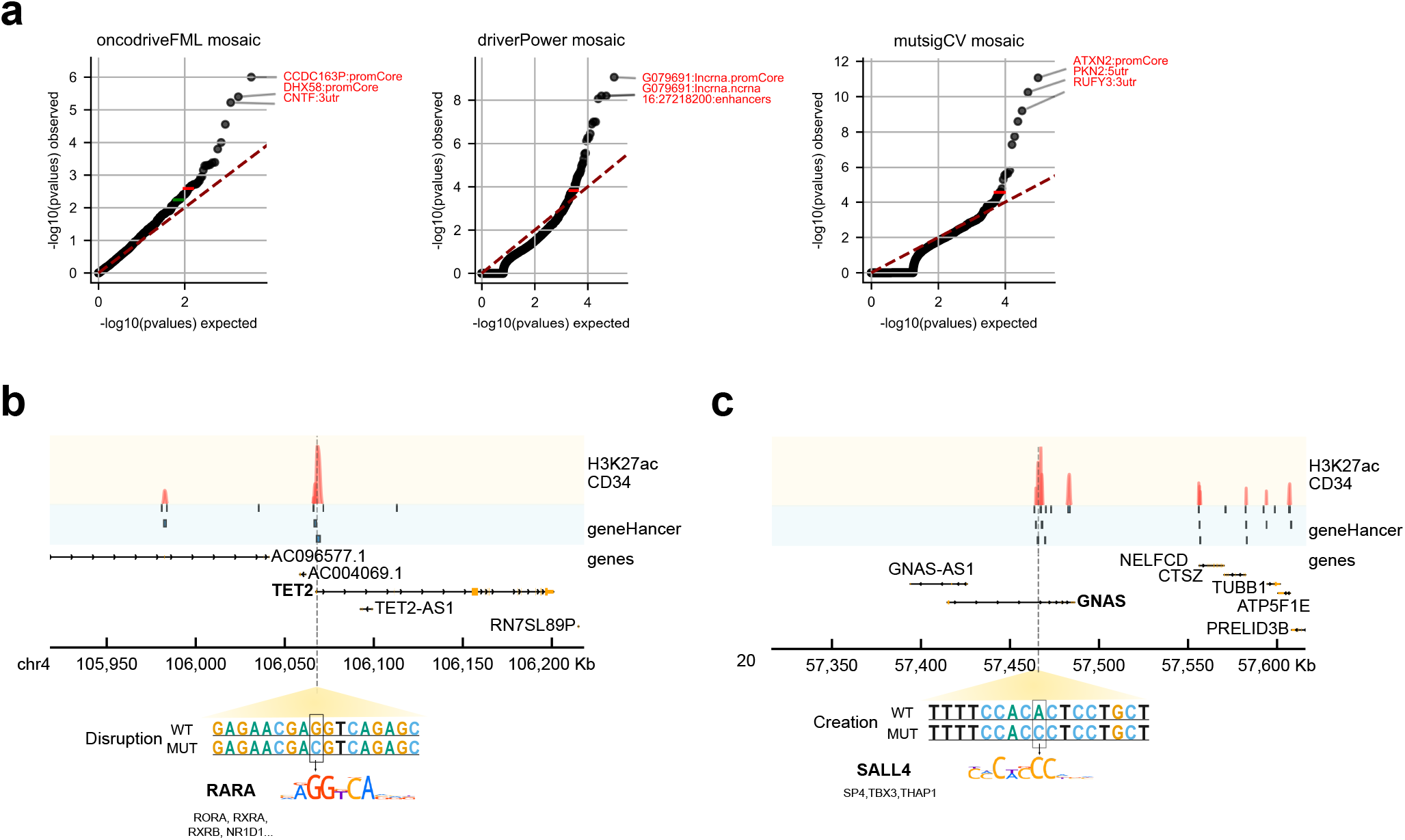
Non-coding mutations in clonal hematopoiesis. (a) Quantile-quantile plots presenting the observed and expected distribution of p-values resulting from the analysis of blood somatic mutations overlapping non-coding genomic elements across the metastasis cohort with three state-of-the-art non-coding driver discovery methods. The names of the most significant non-coding genomic elements are annotated in red in the plot. (b) Example of a mutation that potentially disrupts the binding site of an expressed transcription factor within an enhancer element in the genome of a donor in the metastasis cohort. Enhancers are obtained from a manually curated database (geneHancer) and bear marks of transcriptional activity (i.e, active enhancers) in cells with phenotype close to HSC. Other possible affected transcription factors are also labeled. (c) Example of a mutation that potentially creates a binding site for an expressed transcription factor in one enhancer element. Other possible affected transcription factors are also labeled.

## Supplementary Tables (in accompanying compressed file)

**Table S1. Mutational profiles of signatures active across blood samples in the metastasis cohort**.

**Table S2. Compendium of clonal hematopoiesis driver genes**.

**Table S3. Non-coding elements significant by the analysis of three driver discovery methods**.

**Table S4. Examples of mutations potentially affecting transcription factor binding sites in enhancer elements**.

