## Supplementary Note for "Discovering the drivers of clonal hematopoiesis"

#### Table of Contents

**Comments on the reverse calling** 2

**Identifying genes under positive selection in CH** 2

**Discovery of CH drivers in panel sequencing data** 10

**Significance of the compendium of CH drivers** 12

**References** 14

#### Comments on the reverse calling

The reverse calling approach implemented in this work for the identification of blood somatic mutations has two main advantages with respect to one-sample germline or somatic calling.

First, variants supported by fewer reads than the minimum generally required in a germline calling may be identified. It is this difference that accounts for the gain in sensitivity in the identification of CH across samples in the metastasis cohort exemplified in Figure 1c and Extended Data Fig. 3c. This is a key feature in the aim to repurpose cancer genomics datasets for the discovery of CH driver genes, given that the blood samples across cancer patients cohorts are sequenced at lower depths than their tumor counterparts (e.g., ~40X in the metastasis cohort). Deeper sequencing of these paired blood samples across cancer genomics datasets would definitely favor this repurposing.

Second, the availability of a second sample from the same individual improves the filtering of germline variants of the reverse calling with respect to any filter of polymorphisms implemented posterior to germline calling of the blood sample in isolation. This thus determines that the reverse calling is more specific than a germline calling.

Some other considerations about the reverse calling approach are also worth mentioning. It is possible that the calling algorithm misses some bona-fide CH variants due the homozygous loss of a DNA loci in the tumoral sample. Another issue might be events of loss of heterozygosity in the tumor, which might cause some heterozygous polymorphisms to be detected as a somatic mutation in the algorithm. Nevertheless, this can be tackled with a stringent germline filtering using public polymorphism databases. We have used a very conservative and strict threshold of > 0.0003 AF in gnomAD[1] database to remove putative polymorphisms, as well as run a state-of-the-art machine learning algorithm to distinguish between polymorphisms and somatic mutations[22]. In order to avoid these problems, other non-tumoral control samples, with lower levels of genome instability, might be used.

#### Identifying genes under positive selection in CH

We posit that if somatic mutations identified across healthy blood samples of patients in both cohorts are true hematopoietic mutations that become detectable due to clonal hematopoiesis, then their distribution across the genome is not expected to be completely random. Instead, because clonal hematopoiesis is a phenomenon driven by mutations that provide some hematopoietic stem cells with advantages with respect to others, signals of positive selection are expected to be detectable in their distribution across the genome. Specifically, these signals would be apparent in genes whose mutations are under positive selection in the arisal of clonal hematopoiesis. Thus, we expect that mutations in these CH-related genes exhibit distinct patterns of mutations that deviate from the expected under neutral evolution.

A range of methods to identify these signals of positive selection across genes have been developed in recent years for their application to tumor genomics data with the aim of identifying cancer driver genes. If detected, the signals of positive selection across the somatic mutations in blood samples would thus be a reflection of the clonal expansion triggered by CH.

Therefore, we applied the IntOGen-pipeline[2], which runs seven complementary state-of-the-art methods[3–9] to detect signals of positive selection in the mutation pattern of genes. These methods are designed to detect different deviations of the mutation pattern of genes with respect to their expectation under neutrality --that is, different signals of positive selection. The methods were applied independently to the full and mosaic (see main text) sets of somatic mutations identified across the blood samples of both datasets. As a general rule (i.e., except a couple of cases) the genes with signals of positive selection according to the different methods are significantly enriched for known cancer driver genes --i.e., those that when mutated confer an advantage to somatic cells (CGC[10] genes in Table 1). The same significant enrichment is apparent for known drivers of clonal hematopoiesis (CH in Table 1) and genes that drive specifically myeloid malignancies (Myeloid[11] in Table 1).

We also employed quantile-quantile plots (qqplots) of the results of the different methods (in which the p-values of a set of genes deviate from the uniformity that would be expected under neutrality) to assess their calibration (Fig. 1). Most methods, when run on the mosaic sets of mutations derived from both cohorts exhibit a well calibrated behavior, with only a few significant genes deviating from the diagonal. We hypothesize that in the few cases in which an inflation is observed, the reason is that (unlike in the case of cancer somatic mutations) the sets of mutations employed to run the methods are still contaminated with artifactual mutations with a non-random distribution which may contribute to biasing their results.

Driver identification methods based on different signals of positive selection also show different biases when applied to cancer somatic mutations. This is why we have developed a reasoned approach to combine their outputs that delivers weights on the basis of the perceived credibility of the methods in each cohort[2]. Thus, we anticipate that the biases observed in the results of certain methods when applied to blood somatic mutations in both cohorts, could be solved using this combination approach.

Moreover, because the full set of mutations identified through the reverse calling approach is contaminated by potential sequencing artifacts, we deem the set of clonal hematopoiesis genes identified by the combination of the results of the application of the methods to these filtered catalogs (Mutect and Mosaic) more reliable than that obtained from the full catalog.

In order to derive the list of clonal hematopoiesis drivers, we thus start with genes that are significant (in the combination of methods' outputs) from the analysis of any filtered catalog. Genes that appear significant from the analysis of the full catalog are only included in the final list if they are supported by prior knowledge of their involvement in CH or cancer (i.e., included in the CGC[10]). This way, we generate two lists of putative CH drivers, i.e., one for each cohort. These two lists, directly obtained from the combination algorithm are already very enriched for known CH and cancer genes (Table 2). The genes in these two lists, among which false positives may still be present, are carefully vetted employing criteria that we have developed in a decade-worth of analysis of cancer cohorts (https://intogen.readthedocs.io/en/latest/postprocessing.html#). Specifically, we remove from the lists:

- 10 genes that are not expressed (highest value below 15 fpkms) across a set of HSCs (see methods and Fig. 2)
- 1 gene due to be highly tolerant to Single Nucleotide Polymorphisms[1] (SNP) across human populations
- 2 genes that are frequent false positives of different driver discovery methods
- 1 gene that has more than 3 mutations in one sample, which may be a signal of a local hypermutation process or contamination of germline variants from the reverse calling
- 1 gene that has more than 50% mutations in one of the cohorts associated with COSMIC Signature 9 (https://cancer.sanger.ac.uk/cosmic/signatures), related with the maturation of lymphoid cells

The vetting process yields two lists composed of 26 and 32 genes, all of which are reasonably good candidates of driving clonal hematopoiesis in either cohort (Table 2). The vast majority (23/26 and 26/32) have been mentioned in the literature related to CH, myeloid malignancies, or tumorigenesis in general[12,13]. Although the genes with no prior knowledge of involvement in any of these processes could be bona fide CH genes, further evidence is needed. With the objective to generate a very reliable snapshot of the compendium, these genes are discarded from the compendium, but still listed in Table 2.

**Figure 1. Quantile-quantile plots of driver discovery methods in the detection of CH**. (Figure in next page)
Qqplots of the results of analyzing the mosaic set of mutations are represented. The names of genes that are significant at FDR 0.01, appear in red, while those significant at FDR 0.1, appear in green.

### Figure 1

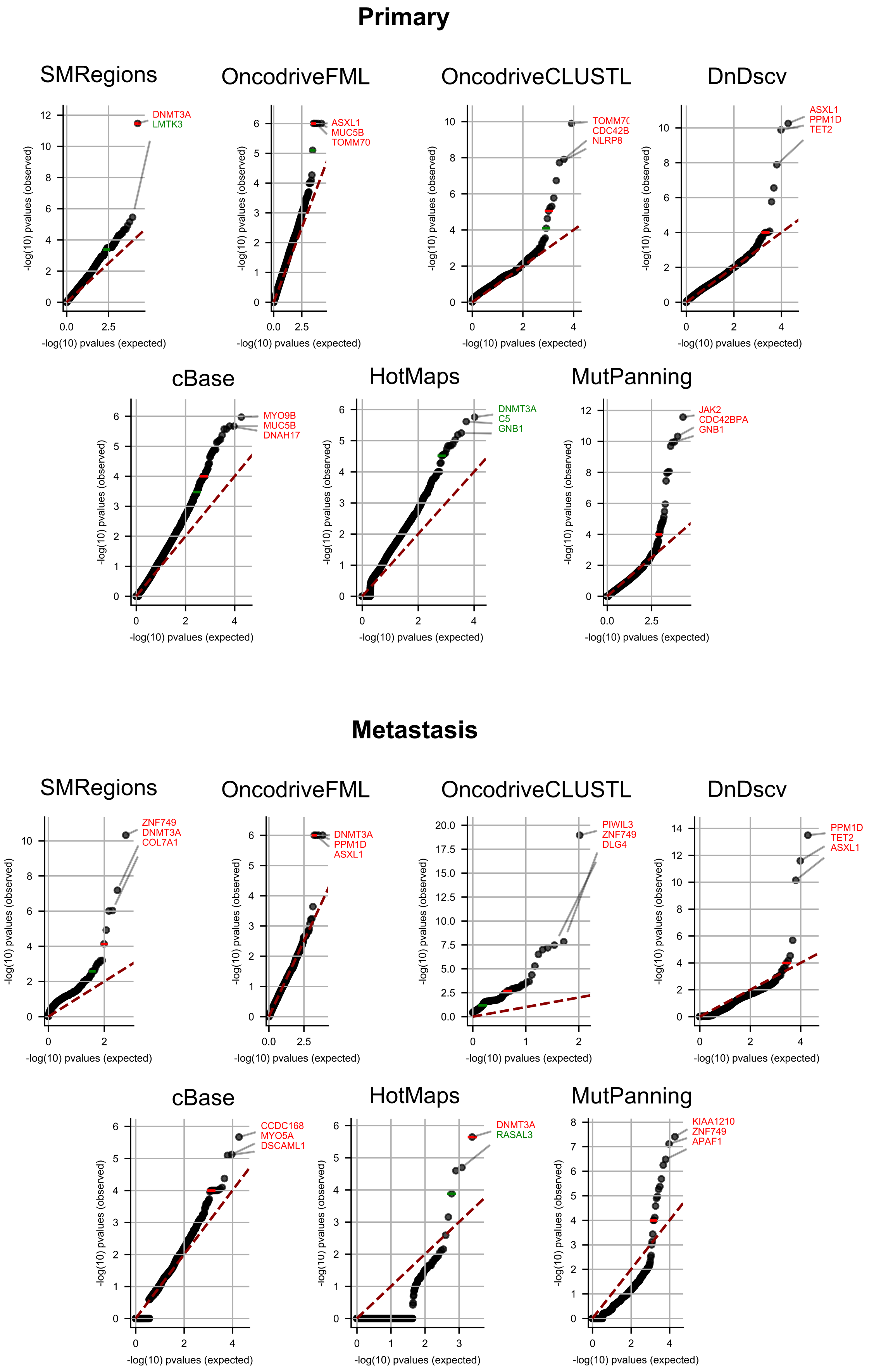

Primary
SMRegions
DNMT3A
LMTK3
OncodriveFML
ASXL1
MUC5B
TOMM70
OncodriveCLUSTL
TOMM7
CDC42B
NLRP8
DnDscv
ASXL1
PPM1D
TET2
cBase
MYO9B
MUC5B
DNAH17
HotMaps
DNMT3A
C5
GNB1
MutPanning
JAK2
CDC42BPA
GNB1
-log(10) pvalues (observed)
-log(10) pvalues (expected)
Metastasis
SMRegions
ZNF749
DNMT3A
COL7A1
OncodriveFML
DNMT3A
PPM1D
ASXL1
OncodriveCLUSTL
PIWIL3
ZNF749
DLG4
DnDscv
PPM1D
TET2
ASXL1
cBase
CCDC168
MYO5A
DSCAML1
HotMaps
DNMT3A
RASAL3
MutPanning
KIAA1210
ZNF749
APAF1

**Figure 2. Expression of genes under positive selection in the primary and metastasis cohorts across HSCs**. (Figure in next page)

Most genes in the list of putative CH drivers identified through the IntOGen pipeline appear expressed across HSCs cells (i.e., above the threshold of 15 fpkm), while only 9 that appear below this value are filtered out. Genes that are filtered out due to lack of expression are marked with '#'; genes that are filtered out due to other criteria (detailed above) bear the '*' label; genes discarded due to lack of literature evidence of involvement in CH or tumorigenesis are labelled '^'. The gene TOMM70, not included in the expression dataset employed for this purpose (see Methods of the main paper) is also excluded from the list of potential CH driver genes.

### Figure 2

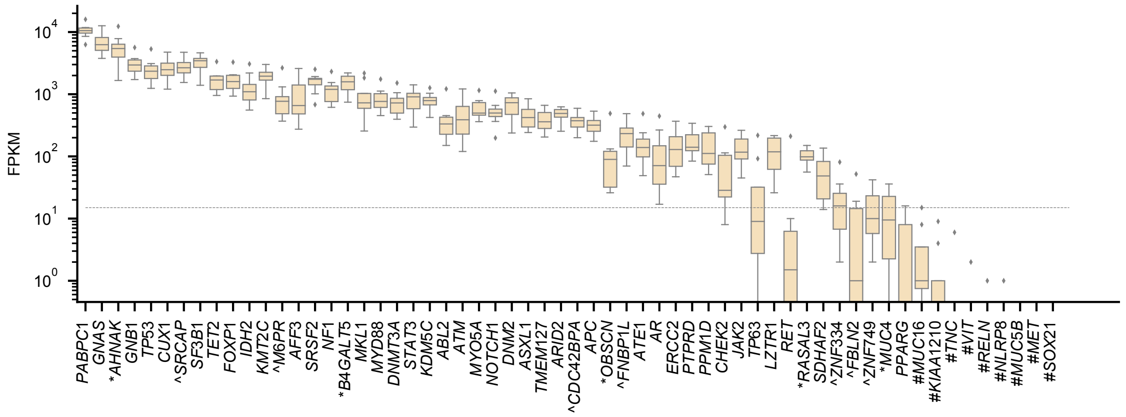

FPKM
$10^4$
$10^3$
$10^2$
$10^1$
$10^0$
PABPC1
GNAS
*AHNAK
GNB1
TP53
CUX1
^SRCAP
SF3B1
TET2
FOXP1
IDH2
KMT2C
^M6PR
AFF3
SRSF2
NF1
*B4GALT5
MKL1
MYD88
DNMT3A
STAT3
KDM5C
ABL2
ATM
MYO5A
NOTCH1
DNM2
ASXL1
TMEM127
ARID2
^CDC42BPA
APC
*OBSCN
^FNBP1L
ATE1
AR
ERCC2
PTPRD
PPM1D
CHEK2
JAK2
TP63
LZTR1
RET
*RASAL3
SDHAF2
^ZNF334
^FBLN2
^ZNF749
*MUC4
PPARG
#MUC16
#KIAA1210
#TNC
#VIT
#RELN
#NLRP8
#MUC5B
#MET
#SOX21

**Table 1. Result of the analysis of blood somatic mutations with driver discovery methods**

| Cohort | Mutations set | Method | Fraction of known genes | | | Total genes | | | Odds-ratio | | | P-value | | |
|---|---|---|---|---|---|---|---|---|---|---|---|---|---|---|
| | | | CH | Myeloid | CGC | Identified by the method | In no list | In any list | CH | Myeloid | CGC | CH | Myeloid | CGC |
| Metastasis | Mutect | Cbase | 0.57 | 0.57 | 0.57 | 7 | 3 | 4 | 2260.4 | 272.1 | 34.4 | 9.4E-12 | 2.2E-08 | 6.3E-05 |
| Metastasis | Mosaic | Cbase | 0.50 | 0.50 | 0.50 | 8 | 4 | 4 | 1695.2 | 204.0 | 25.8 | 1.9E-11 | 4.3E-08 | 1.2E-04 |
| Metastasis | Full | Cbase | 0.57 | 0.57 | 0.57 | 7 | 3 | 4 | 2260.4 | 272.1 | 34.4 | 9.4E-12 | 2.2E-08 | 6.3E-05 |
| Primary | Mosaic | Cbase | 0.27 | 0.27 | 0.27 | 15 | 11 | 4 | 616.2 | 74.2 | 9.4 | 3.7E-10 | 8.2E-07 | 1.9E-03 |
| Primary | Full | Cbase | 0.27 | 0.27 | 0.33 | 15 | 10 | 5 | 616.2 | 74.2 | 12.9 | 3.7E-10 | 8.2E-07 | 1.6E-04 |
| Metastasis | Mutect | OncodriveCLUSTL | 1.00 | 1.00 | 1.00 | 1 | 0 | 1 | NA | NA | NA | NA | NA | NA |
| Metastasis | Mosaic | OncodriveCLUSTL | 0.17 | 0.17 | 0.17 | 23 | 19 | 4 | 16.8 | NA | 8.3 | 8.2E-03 | 1.9E-03 | 2.1E-02 |
| Metastasis | Full | OncodriveCLUSTL | 0.02 | 0.04 | 0.06 | 130 | 122 | 8 | 2.0 | 2.0 | 0.7 | 3.9E-01 | 2.0E-01 | 4.8E-01 |
| Primary | Mosaic | OncodriveCLUSTL | 0.13 | 0.13 | 0.13 | 8 | 7 | 1 | 107.8 | 21.8 | 2.8 | 1.1E-02 | 5.2E-02 | 3.3E-01 |
| Primary | Full | OncodriveCLUSTL | 0.05 | 0.05 | 0.11 | 66 | 59 | 7 | 68.1 | 8.1 | 2.7 | 2.7E-05 | 7.3E-03 | 2.3E-02 |
| Metastasis | Mutect | SMRegions | 1.00 | 1.00 | 1.00 | 1 | 0 | 1 | NA | NA | NA | NA | NA | NA |
| Metastasis | Mosaic | SMRegions | 0.33 | 0.33 | 0.33 | 6 | 4 | 2 | 52.8 | 19.5 | 4.7 | 2.8E-03 | 1.2E-02 | 1.2E-01 |
| Primary | Mosaic | SMRegions | 1.00 | 1.00 | 1.00 | 1 | 0 | 1 | NA | NA | NA | NA | NA | NA |
| Metastasis | Full | SMRegions | 0.08 | 0.08 | 0.08 | 37 | 33 | 4 | 23.3 | 6.6 | 1.2 | 8.3E-04 | 1.6E-02 | 7.4E-01 |
| Primary | Full | SMRegions | 0.25 | 0.19 | 0.19 | 16 | 12 | 4 | 539.0 | 37.6 | 4.8 | 8.7E-10 | 1.3E-04 | 3.4E-02 |
| Metastasis | Mosaic | DnDsCV | 1.00 | 1.00 | 1.00 | 4 | 0 | 4 | NA | NA | NA | 2.4E-13 | 5.6E-10 | 1.8E-06 |
| Metastasis | Mutect | DnDsCV | 0.09 | 0.09 | 0.14 | 43 | 37 | 6 | 178.9 | 21.5 | 4.3 | 2.9E-08 | 5.9E-05 | 4.5E-03 |
| Metastasis | Full | DnDsCV | 0.80 | 0.80 | 0.80 | 5 | 1 | 4 | 6992.7 | 841.8 | 106.1 | 1.2E-12 | 2.8E-09 | 8.6E-06 |
| Primary | Mosaic | DnDsCV | 0.50 | 0.50 | 0.50 | 8 | 4 | 4 | 1747.9 | 210.4 | 26.5 | 1.7E-11 | 3.8E-08 | 1.1E-04 |
| Primary | Full | DnDsCV | 0.17 | 0.17 | 0.17 | 24 | 20 | 4 | 349.3 | 42.0 | 5.3 | 2.5E-09 | 5.5E-06 | 1.0E-02 |
| Metastasis | Mutect | OncodriveFML | 1.00 | 1.00 | 1.00 | 3 | 0 | 3 | NA | NA | NA | 8.3E-07 | 5.5E-06 | 3.1E-04 |
| Metastasis | Mosaic | OncodriveFML | 1.00 | 1.00 | 1.00 | 4 | 0 | 4 | NA | NA | NA | 1.6E-10 | 2.2E-08 | 2.1E-05 |
| Metastasis | Full | OncodriveFML | 0.33 | 0.33 | 0.42 | 12 | 7 | 5 | 393.1 | 65.1 | 12.9 | 3.3E-09 | 1.9E-06 | 2.4E-04 |
| Primary | Mosaic | OncodriveFML | 0.60 | 0.60 | 0.60 | 5 | 2 | 3 | 2062.0 | 289.8 | 36.8 | 6.1E-09 | 1.5E-06 | 5.7E-04 |
| Primary | Full | OncodriveFML | 0.36 | 0.36 | 0.36 | 11 | 7 | 4 | 904.8 | 115.2 | 14.4 | 1.2E-10 | 2.1E-07 | 5.7E-04 |
| Metastasis | Mosaic | Mutpanning | 0.455 | 0.455 | 0.455 | 11 | 6 | 5 | 1551.2 | 171.6 | 21.5 | 7.4E-14 | 1.4E-09 | 2.8E-05 |
| Metastasis | Full | Mutpanning | 0.192 | 0.192 | 0.269 | 26 | 19 | 7 | 442.1 | 48.9 | 9.5 | 1.1E-11 | 1.9E-07 | 3.6E-05 |
| Primary | Mosaic | Mutpanning | 0.467 | 0.400 | 0.400 | 15 | 8 | 7 | 2036.0 | 138.8 | 17.2 | 2.7E-19 | 7.2E-11 | 1.0E-05 |
| Primary | Full | Mutpanning | 0.125 | 0.111 | 0.139 | 72 | 61 | 11 | 441.4 | 26.5 | 4.2 | 5.7E-19 | 3.1E-09 | 3.5E-04 |
| Metastasis | Mosaic | HotMaps | 1.000 | 1.000 | 1.000 | 1 | 0 | 1 | NA | NA | NA | NA | NA | NA |
| Metastasis | Full | HotMaps | 0.250 | 0.250 | 0.250 | 4 | 3 | 1 | 144.7 | 37.7 | 5.7 | 1.0E-02 | 3.5E-02 | 2.0E-01 |

**Table 2. List of genes with signals of positive selection across the primary and metastasis cohort after removal of known artifacts.**

| COHORT | SET | GENE | DECISION |
|---|---|---|---|
| Primary | Mosaic | ABL2 | IN COMPENDIUM |
| Primary | Full | AFF3 | IN COMPENDIUM |
| Primary | Mosaic | AFF3 | IN COMPENDIUM |
| Metastasis | Mutect | APC | IN COMPENDIUM |
| Primary | Full | AR | IN COMPENDIUM |
| Metastasis | Mosaic | ARID2 | IN COMPENDIUM |
| Metastasis | Mutect | ARID2 | IN COMPENDIUM |
| Metastasis | Full | ASXL1 | IN COMPENDIUM |
| Metastasis | Mosaic | ASXL1 | IN COMPENDIUM |
| Metastasis | Mutect | ASXL1 | IN COMPENDIUM |
| Primary | Full | ASXL1 | IN COMPENDIUM |
| Primary | Mosaic | ASXL1 | IN COMPENDIUM |
| Metastasis | Mosaic | ATE1 | IN COMPENDIUM |
| Metastasis | Full | ATM | IN COMPENDIUM |
| Metastasis | Mosaic | ATM | IN COMPENDIUM |
| Metastasis | Mutect | ATM | IN COMPENDIUM |
| Primary | Mosaic | ATM | IN COMPENDIUM |
| Metastasis | Mosaic | CHEK2 | IN COMPENDIUM |
| Primary | Mosaic | CHEK2 | IN COMPENDIUM |
| Metastasis | Mutect | CUX1 | IN COMPENDIUM |
| Metastasis | Full | DNM2 | IN COMPENDIUM |
| Metastasis | Full | DNMT3A | IN COMPENDIUM |
| Metastasis | Mosaic | DNMT3A | IN COMPENDIUM |
| Metastasis | Mutect | DNMT3A | IN COMPENDIUM |
| Primary | Full | DNMT3A | IN COMPENDIUM |
| Primary | Mosaic | DNMT3A | IN COMPENDIUM |
| Primary | Mosaic | ERCC2 | IN COMPENDIUM |
| Metastasis | Mosaic | FOXP1 | IN COMPENDIUM |
| Primary | Full | GNAS | IN COMPENDIUM |
| Primary | Mosaic | GNAS | IN COMPENDIUM |
| Metastasis | Full | GNB1 | IN COMPENDIUM |
| Primary | Full | GNB1 | IN COMPENDIUM |
| Primary | Mosaic | GNB1 | IN COMPENDIUM |
| Metastasis | Full | IDH2 | IN COMPENDIUM |
| Metastasis | Full | JAK2 | IN COMPENDIUM |
| Metastasis | Mosaic | JAK2 | IN COMPENDIUM |
| Primary | Full | JAK2 | IN COMPENDIUM |
| Primary | Mosaic | JAK2 | IN COMPENDIUM |
| Metastasis | Full | KDM5C | IN COMPENDIUM |
| Primary | Mosaic | KMT2C | IN COMPENDIUM |
| Primary | Mosaic | LZTR1 | IN COMPENDIUM |
| Metastasis | Full | MKL1 | IN COMPENDIUM |
| Primary | Full | MYD88 | IN COMPENDIUM |
| Metastasis | Full | MYO5A | IN COMPENDIUM |
| Metastasis | Mosaic | MYO5A | IN COMPENDIUM |
| Metastasis | Mosaic | NF1 | IN COMPENDIUM |
| Metastasis | Full | NOTCH1 | IN COMPENDIUM |
| Primary | Full | PABPC1 | IN COMPENDIUM |
| Primary | Mosaic | PPARG | IN COMPENDIUM |
| Metastasis | Full | PPM1D | IN COMPENDIUM |
| Metastasis | Mosaic | PPM1D | IN COMPENDIUM |
| Metastasis | Mutect | PPM1D | IN COMPENDIUM |
| Primary | Full | PPM1D | IN COMPENDIUM |
| Primary | Mosaic | PPM1D | IN COMPENDIUM |
| Metastasis | Mosaic | PTPRD | IN COMPENDIUM |
| Metastasis | Mosaic | RET | IN COMPENDIUM |
| Primary | Full | SDHAF2 | IN COMPENDIUM |
| Metastasis | Mosaic | SF3B1 | IN COMPENDIUM |
| Metastasis | Mutect | SF3B1 | IN COMPENDIUM |
| Metastasis | Full | SRSF2 | IN COMPENDIUM |
| Metastasis | Mutect | SRSF2 | IN COMPENDIUM |
| Primary | Full | SRSF2 | IN COMPENDIUM |
| Primary | Mosaic | SRSF2 | IN COMPENDIUM |
| Primary | Full | STAT3 | IN COMPENDIUM |
| Metastasis | Full | TET2 | IN COMPENDIUM |
| Metastasis | Mosaic | TET2 | IN COMPENDIUM |
| Metastasis | Mutect | TET2 | IN COMPENDIUM |
| Primary | Full | TET2 | IN COMPENDIUM |
| Primary | Mosaic | TET2 | IN COMPENDIUM |
| Primary | Full | TMEM127 | IN COMPENDIUM |
| Metastasis | Full | TP53 | IN COMPENDIUM |
| Metastasis | Mosaic | TP53 | IN COMPENDIUM |
| Metastasis | Mutect | TP53 | IN COMPENDIUM |
| Primary | Full | TP53 | IN COMPENDIUM |
| Primary | Mosaic | TP53 | IN COMPENDIUM |
| Metastasis | Mosaic | TP63 | IN COMPENDIUM |
| Primary | Mosaic | B4GALT5 | DISCARDED |
| Primary | Full | CDC42BPA | DISCARDED |
| Primary | Mosaic | CDC42BPA | DISCARDED |
| Metastasis | Full | FBLN2 | DISCARDED |
| Metastasis | Mosaic | FBLN2 | DISCARDED |
| Metastasis | Mosaic | FNBP1L | DISCARDED |
| Primary | Full | M6PR | DISCARDED |
| Primary | Mosaic | M6PR | DISCARDED |
| Metastasis | Mosaic | RASAL3 | DISCARDED |
| Metastasis | Mosaic | SRCAP | DISCARDED |
| Metastasis | Mutect | SRCAP | DISCARDED |
| Metastasis | Full | ZNF334 | DISCARDED |
| Metastasis | Mosaic | ZNF334 | DISCARDED |
| Metastasis | Full | ZNF749 | DISCARDED |
| Metastasis | Mosaic | ZNF749 | DISCARDED |

#### Discovery of CH drivers in panel sequencing data

In many clinically oriented initiatives, a subset of all protein-coding genes in solid tumors has been sequenced. In some cases, such as the MSK-IMPACT[14,15], a paired blood sample has also been sequenced with the aim of correctly calling tumor somatic mutations. In two recent studies, the germline variants identified across 24,146 such blood samples have been filtered with those appearing in the tumor sample from the same patient, thus yielding likely blood somatic mutations[11,16].

We hypothesized that the same rationale of discovery of clonal hematopoiesis driver genes presented here could be applied to this cohort (targeted cohort). The genes in the MSK-IMPACT panel have been selected because they are involved in tumor development; most are included in the CGC. In the two aforementioned studies, blood somatic mutations affecting them have been taken as evidence of clonal hematopoiesis. We propose that identifying signals of positive selection in this cohort would fulfill at least two main objectives. First, due to the number of samples included in the cohort, and thus the high statistical power, the discovery would most likely extend the list of drivers of clonal hematopoiesis. Second, given the nature of the genes included in the panel, it would identify cancer driver genes that do not show any evidence to be drivers of clonal hematopoiesis, and thus identified blood mutations may be passenger mutations.

Unlike the discovery described in the paper for the metastasis and the primary cohorts, the “panel discovery” of CH drivers is limited to the 468 genes included in the MSK-IMPACT panel. Furthermore, only driver discovery methods that rely on local background models --i.e., capable of computing a background model from the fragment of the genome covered by the panel-- could be employed. We thus applied OncodriveFML[3], OncodriveCLUSTL[5], DnDscv[4] (without the mutation rate covariates, as described in ref[17]), and SMRegions[6] to the blood somatic mutations identified in these 468 genes across 24,146 samples.

Forty-four drivers of clonal hematopoiesis are identified by at least one method in the targeted cohort discovery (Fig. 3a). Twenty-eight panel CH drivers are identified by more than one method. Twenty-nine of the panel CH driver genes are only identified across the targeted cohort (Fig. 2d of the main paper). Interestingly, 9 genes included in the panel are not identified as CH drivers by the targeted cohort discovery, but are identified either in the primary (4) or in the metastasis (5) cohorts (Fig. 3b). This supports the notion that an extensive effort, including new cohorts is the path to the discovery of the compendium of drivers of clonal hematopoiesis, as has been demonstrated in cancer[2].

One of the main outcomes of this work is an unbiased snapshot of the compendium of clonal hematopoiesis driver genes, presented in Table S2 of the main paper and available at www.intogen.org/ch. This compendium is integrated by the genes that exhibit signals of positive selection in their mutational patterns in at least one of the three cohorts analyzed in the paper (whole-exome primary, whole-genome metastasis, and targeted).

### Figure 3

**a**

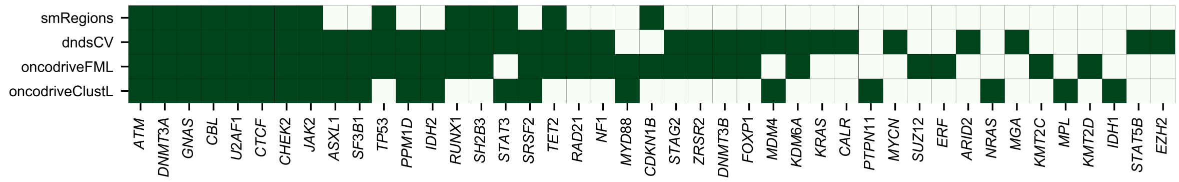
smRegions
dndsCV
oncodriveFML
oncodriveClustL
ATM
DNMT3A
GNAS
CBL
U2AF1
CTCF
CHEK2
JAK2
ASXL1
SF3B1
TP53
PPM1D
IDH2
RUNX1
SH2B3
STAT3
SRSF2
TET2
RAD21
NF1
MYD88
CDKN1B
STAG2
ZRSR2
DNMT3B
FOXP1
MDM4
KDM6A
KRAS
CALR
PTPN11
MYCN
SUZ12
ERF
ARID2
NRAS
MGA
KMT2C
MPL
KMT2D
IDH1
STAT5B
EZH2

**b**

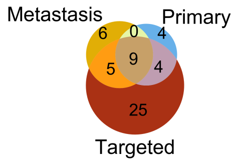
Metastasis
Primary
6
0
4
9
5
4
25
Targeted

**Figure 3. Clonal hematopoiesis drivers identified across targeted sequenced blood samples.**
(a) Genes with signals of positive selection (identified by 4 methods as shown in the heatmap) across blood samples probed with the MSK-IMPACT panel (targeted cohort).
(b) Some genes included in the MSK-IMPACT panel only exhibit signals of positive selection in the Primary or Metastasis cohorts (represented by the Venn diagram). This illustrates the importance of carrying out a discovery of CH drivers across cohorts.

Interestingly, although genes in the MSK-IMPACT panels have been selected due to their involvement in tumorigenesis, most of them (424) show no signals of positive selection across these 24,146 blood samples. This includes some genes that appear recurrently mutated in the cohort. For example, EGFR or MED12 well known cancer genes mutated in 58 and 45 blood samples (respectively) in the targeted cohort do not show any signal of positive selection in their mutation patterns. These mutations therefore do not have any support of being drivers of clonal hematopoiesis, and the mutations detected in blood might just be passenger mutations

This observation has important implications for the detection of CH through the identification of somatic mutations in these genes across blood samples. Traditionally, the occurrence of CH in a blood sample is detected either through a mutation affecting a CH driver, or because a number of hematopoiesis mutations are identified in the sample (both approaches are illustrated in Fig. 4g of the main paper). The identification of a mutation in a cancer driver gene which is not a CH driver (in the absence of a critical mass of hematopoiesis mutations detected in the same sample) may lead to a spurious classification of the sample as CH.

#### Significance of the compendium of CH drivers

As pointed out in the Discussion of the main paper, the availability of the compendium of clonal hematopoiesis driver genes is significant in at least two regards. First, the compendium of CH drivers will help advance the research on the molecular mechanisms underlying clonal hematopoiesis faced with different evolutionary constraints (cytotoxic treatments, tobacco carcinogens, etc). Second, knowing the compendium of CH drivers will improve the diagnosis of the condition across human donors, by helping distinguish mutations that more likely drive CH in a donor's blood sample. One can easily imagine that the completion of the compendium will lead to the development of targeted sequencing panels focused on CH drivers. These would guarantee sequencing relevant genes --and, eventually other genomic elements-- at higher depth, thus discovering the condition as early as possible in population screenings.

A second step to take in the direction of identifying CH-driving mutations remains. As the study of tumorigenesis has revealed, not all mutations in CH driver genes will be equally capable of driving clonal hematopoiesis. This is already apparent in the distribution of observed mutations along the sequence of CH drivers presented in Figure 3 of the main paper. The example of mutations affecting PPM1D is very eloquent in this sense. In clonal hematopoiesis cases, mutations in this gene tend to be truncating, resulting in the loss of a degron located in the C-terminal portion of the protein, thus leading to its abnormal stabilization which results in decreased levels of the active TP53 protein product[18–21].
